# Luminal transport through intact endoplasmic reticulum limits the magnitude of localized Ca^2+^ signals

**DOI:** 10.1101/2023.06.23.546357

**Authors:** Cécile C. Crapart, Zubenelgenubi C. Scott, Tasuku Konno, Aman Sharma, Pierre Parutto, David M. D. Bailey, Laura M. Westrate, Edward Avezov, Elena F. Koslover

**Author notes:** To whom correspondence should be addressed. E-mail: ea347cam.ac.uk (EA), ekosloverucsd.edu (EFK). Please provide details of author contributions here. The authors declare no competing interest.

## Abstract

The endoplasmic reticulum (ER) forms an interconnected network of tubules stretching throughout the cell. Understanding how ER functionality relies on its structural organization is crucial for elucidating cellular vulnerability to ER perturbations, which have been implicated in several neuronal pathologies. One of the key functions of the ER is enabling Ca^2+^ signalling by storing large quantities of this ion and releasing it into the cytoplasm in a spatiotemporally controlled manner. Through a combination of physical modeling and livecell imaging, we demonstrate that alterations in ER shape significantly impact its ability to support efficient local Ca^2+^ releases, due to hindered transport of luminal content within the ER. Our model reveals that rapid Ca^2+^ release necessitates mobile luminal buffer proteins with moderate binding strength, moving through a well-connected network of ER tubules. These findings provide insight into the functional advantages of normal ER architecture, emphasizing its importance as a kinetically efficient intracellular Ca^2+^ delivery system.

**Significance Statement:** The peripheral endoplasmic reticulum forms a continuous network of tubules extending through the entire cell. One of the key functional roles of the ER is the release of Ca^2+^ ions into the cytosol to support a broad diversity of intracellular signaling processes. Such release events are enabled by the high Ca^2+^ storage capacity of the ER. This work demonstrates that mobile Ca^2+^binding buffer proteins and a well-connected lattice-like architecture of the ER network are optimal to supply local Ca^2+^ signals and that changes in ER structure can modulate Ca^2+^ release. By linking transport kinetics to Ca^2+^ release, we demonstrate a key functional role for the interconnected network architecture of the ER.

---

The endoplasmic reticulum (ER) is a membrane-enclosed organelle with characteristic structural subdomains, including stacks of membranous sheets contiguously interconnected with a network of tubules, all sharing a common lumen. The balance between tubules, junctions, and sheets varies substantially among cell types and conditions, and is thought to be controlled by a combination of ER-shaping morphogen proteins and tension in the ER membrane (1–4). ER sheet morphologies convey a clear functional advantage by maximizing surface area to accommodate ribosomes for secretory protein synthesis (5, 6). However, the functional role of the highly looped interconnected architecture of the peripheral ER network remains obscure. This knowledge gap hinders our understanding of the diseases associated with ER shaping proteins. In particular, mutations in ER morphogens cause Hereditary Spastic Paraplegia and ALS (motor neuron diseases) (7) and their protein level alterations are associated with Alzheimer’s disease (8, 9). The apparent selective vulnerability of neuronal cells, with their extensive ER tubule-containing periphery, has not been mechanistically rationalized. The normally shaped and functioning cell-wide network of ER tubules is thought to allow delivery of bioactive molecules, including proteins, lipids and Ca^2+^ ions, for signalling or supply to other organelles (10, 11). However, it remains unclear how the morphology of the ER modulates its function as an intracellular transport network, particularly its role in the delivery of Ca^2+^ for signalling events. ER-facilitated Ca^2+^ signaling enables selective regulation of diverse processes in numerous cell types: local Ca^2+^ transients (“puffs”, originally detected in Xenopus oocytes during maturation (13, 14)) are detectable in cultured cell lines (eg: HeLa (15), SHSY5Y (16)), pancreatic acinar cells accompanying secretion, (17) and smooth muscle cells, governing contraction/relaxation (18, 19). Localized Ca^2+^ transients and microdomains also play critical roles in healthy neuronal and astrocytic signaling (20–24).

The role of the ER in Ca^2+^ signaling is facilitated by its Ca^2+^ storage capacity: the narrow ER lumen holds free and protein buffer-bound Ca^2+^ at concentrations three orders of magnitude higher than the cytoplasm. The ions can be released from the ER on demand through the opening of inositol 1,4,5-triphosphate receptor-channels (IP_3_Rs) (15, 25) or ryanodine receptors (RyRs) (26). Other cellular ions can influence ER Ca^2+^ fluxes. Potassium (K^+^) ion flows through large-conductance channels help modulate electrochemical potential changes across the ER membrane during Ca^2+^ release (27, 28). Potassium-hydrogen exhangers and small conductance calcium-sensitive K^+^ channels work together with chloride channels to maintain electroneutrality during Ca^2+^ uptake by the ER (29). Elevated cytosolic levels of zinc ions (Zn^2+^) can induce Ca^2+^ release from the ER, and alterations in ER Ca2+ levels reciprocally influence ER Zn^2+^ uptake (30).

Channel opening can generate Ca^2+^ responses of various extent, ranging from elementary local Ca^2+^ signals upon opening of a single or a few IP_3_Rs (31, 32), to large-scale events that can propagate throughout the cell as a saltatory wave via Ca^2+^-induced Ca^2+^ release (33, 34), triggering whole-cell response (35–37). Localized Ca^2+^ release events, in particular, underscore the need to elucidate how the architecture of interconnected ER tubules may help support Ca^2+^ signaling dynamics by tunneling ions through the ER lumen. These elementary local events form the topic of the current study.

Modeling studies of Ca^2+^ release have highlighted the importance of several key physical parameters, including the distribution and opening probability of channels on the ER membrane (38–41), the buffering capacity of luminal Ca^2+^binding proteins (38), and the relative Ca^2+^ diffusivity in the lumen and cytoplasm (42). Model results have indicated that the mobility of luminal buffer plays an important role in supporting Ca^2+^ release currents when free Ca^2+^ diffusivity in the lumen is much less than the cytoplasm (38), and that the luminal buffering capacity largely determines the total Ca^2+^ levels stored in the lumen (39).

Crucially, ER Ca^2+^ signalling requires enough free ions available to steeply raise the local cytosolic Ca^2+^ levels (43). The release of Ca^2+^ ions must work against several offsetting effects: strong dilution due to their rapid diffusion in the cytoplasm (44), cytoplasmic buffering (45), and active clearing/reuptake (46). This raises the question of whether the ER can sustain an adequate supply of Ca^2+^ for release by drawing exclusively on its local pool, or whether local release events require tunnelling of the ion from more distant ER regions. In the latter case, we expect the Ca^2+^ signalling activity to be dependent on characteristics that determine the efficiency of transport and mixing of luminal content, including network connectivity and the kinetic parameters of motion through the ER lumen. Since the vast majority of ER luminal Ca^2+^ is bound to abundant high-capacity buffer proteins (e.g. calreticulin, with 25 Ca^2+^ binding sites (47)), the mobility and binding affinity of the carrier proteins may also serve as potentially important parameters for optimizing Ca^2+^ release efficiency.

Intra-luminal Ca^2+^ gradients that can arise during release have been noted in models on both bulk geometries (38), and realistic extracted ER architectures (39). However, the interplay between changes to ER morphology, often observed in (patho)physiological circumstances and alterations in Ca^2+^ release dynamics remain unexplored. Here, we seek to investigate how the interconnected morphology of the peripheral ER tubular network helps support the transport and release of buffered Ca^2+^ from the lumen.

To gain a quantitative understanding of the impact of intra-ER luminal transport on localized Ca^2+^ release, we construct a coarse-grained physical model that incorporates localized permeability of the ER membrane to Ca^2+^ ions, equilibrated binding of luminal Ca^2+^ to buffer sites, and diffusive and advective transport through a tubular network geometry. The model reveals that rapid Ca^2+^ release requires mobile buffer proteins with a relatively low binding strength, transported through a well-connected ER network lattice. Further, to validate the predictions of the *in silico* model, we have established a set of tools to perturb various aspects of ER structure via morphogen manipulation, leading to partial luminal fragmentation through tubule narrowing or increase in tubule length through junction depletion. Using all-optical imaging of Ca^2+^ dynamics and fluorescence lifetime imaging (FLIM) measurements of Ca^2+^ concentrations in absolute terms allows exploration of how Ca^2+^ handling in the ER depends on its modified morphology. The resulting experimental data confirm that morphological changes elicited by ER-shaping gene manipulations result in slower luminal material transport and consequently smaller local Ca^2+^ releases. These results provide a functional role for the complex architecture of the ER network as an intracellular Ca^2+^ transport system. The conceptual framework presented here can help establish the optimal ER structure for various physiological conditions and rationalize pathologies associated with ER perturbations.

## Results

### Purely local Ca^2+^ release is insufficient to raise cytoplasmic levels

To explore how the Ca^2+^ release function of the ER is dependent on its structural features, we set out to model the various physical effects that govern local Ca^2+^ currents out of the lumen. We begin with a maximally simple aspatial model that incorporates equilibrated luminal buffer binding, an effective coarse-grained permeability for Ca^2+^ through the ER membrane, and dilution upon cytoplasmic release. In this initial model, the intra-ER transport is not included. Our goal is to establish the extent to which spatiotemporal elevation of cytoplasmic Ca^2+^ can be achieved by the locally available pool in the ER lumen. The results in this section mirror past models of Ca^2+^ release currents (38) and provide a standard of comparison to the subsequent modeling of explicit geometries. Further, these results point to the insufficiency of purely local release for generating observed magnitudes of cytoplasmic Ca^2+^ elevation.

In the purely local model, an isolated region of ER (radius *R* = 0.25*μ*m) maintains a well-mixed lumen and releases Ca^2+^ into the cytoplasm with permeability *P* (Fig. 1a). The effective permeability parameter (units of *μ*m/s) subsumes details of individual channel densities and opening probabilities (48). Our focus is on the release process itself, and we do not include various Ca^2+^ reuptake and clearance mechanisms that would only serve to lower the cytoplasmic Ca^2+^ levels.

**Fig. 1.**
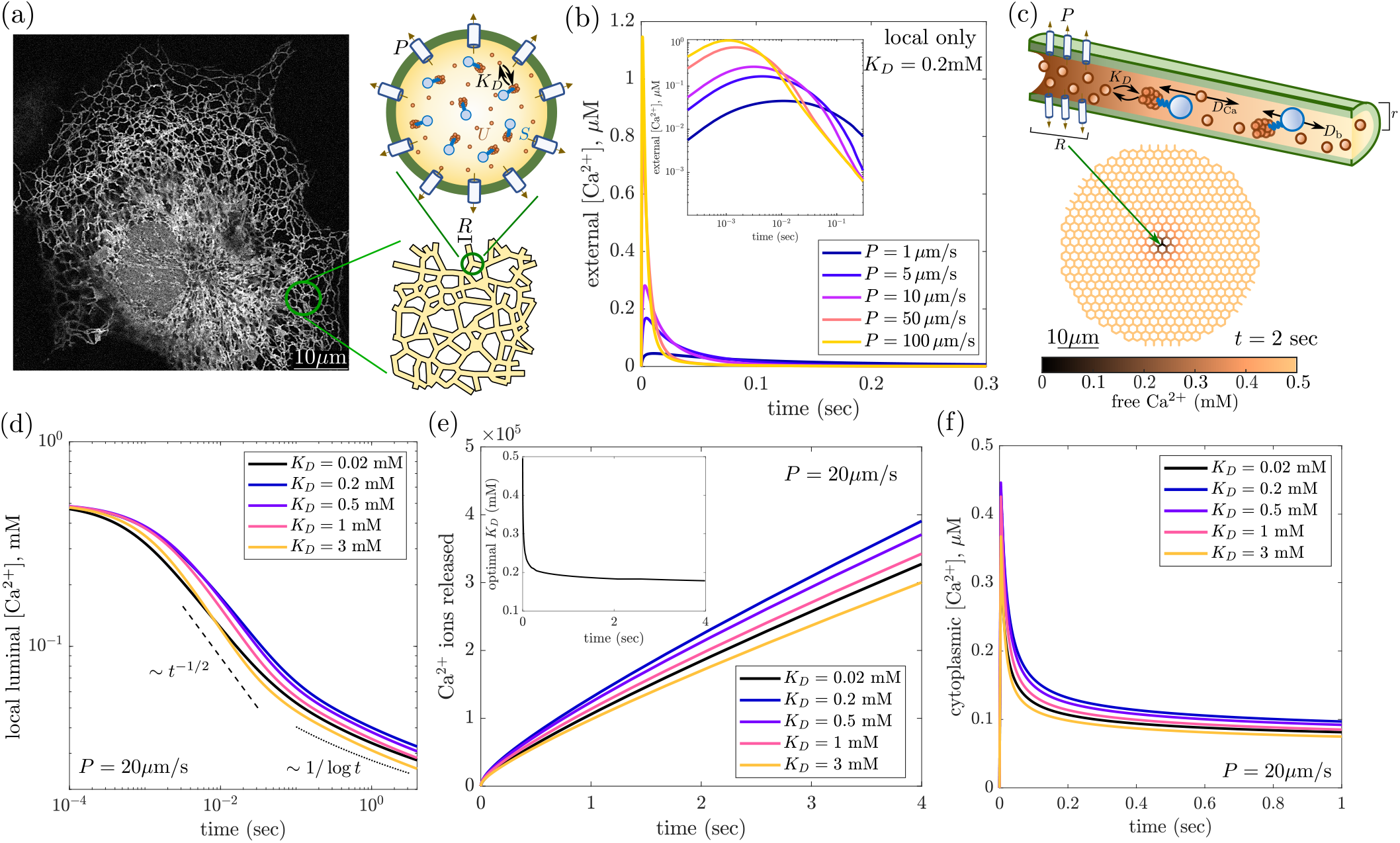
Quantitative modeling demonstrates that Ca^2+^ release requires transport through tubular network and depends on buffer strength. (a) Sample image of peripheral ER network in COS-7 cell. Initial model represents local Ca^2+^ releasing region as a well-mixed domain. (b) Cytoplasmic Ca^2+^ levels predicted from well-mixed model, showing very low local Ca^2+^ event magnitudes from release of only locally available Ca^2+^, for different values of permeability *P*. Inset shows same curves on log-log axes. (c) Snapshot of spatially resolved model for Ca^2+^ release from a honeycomb network structure. Color corresponds to free luminal Ca^2+^. (d) From spatially-resolved model, average free [Ca^2+^]_ER_ in local release region plotted over time for different values of buffer binding strength *K*_*D*_. Dashed line shows short-time scaling expected for 1D delivery through tubules. Dotted line shows long-time scaling expected for 2D delivery. (e) Cumulative amount of Ca^2+^ ions released from the model network over time, for different values of *K*_*D*_ (legend as in d). Inset shows optimal *K*_*D*_ value for maximum cumulative Ca^2+^ released as a function of time. (f) Cytoplasmic Ca^2+^ concentrations after release from model network, averaged over a sphere of *R*_loc_. Default parameter values are *P* = 20*μ*m/s, *S* = 2.5mM, *R* = 0.25*μ*m. Cytoplasmic spread assumes *D*_cyto_ = 200*μ*m^2^ */*s, *R*_loc_ = 1*μ*m.

The cytoplasm is treated as a bulk three-dimensional continuum. The release region contains a length *L* of ER tubules with radius *r*. The scaling of tubule length with the size of the release region depends on the architecture of the network. Here we consider a small region surrounding a degree-3 junction node, which yields an estimate of *L ≈* 3*R*. Given the narrow width of ER tubules, the ratio of ER lumen to cytoplasmic volume in a spherical region surrounding the local release site is very small: *V*_ER_*/V*_cyto_ *≈* 3*r*^2^*L/*4*R*^3^ *≈* 0.09. Furthermore, over time the released Ca^2+^ will spread out through the three-dimensional cytoplasm, implying that local cytoplasmic Ca^2+^ elevation must be quite limited despite the high concentration of ER luminal Ca^2+^. Such dilution effects are known to underlie the relatively low elevation of cytoplasmic Ca^2+^ as compared to luminal Ca^2+^ levels (49). Binding to cytosplasmic buffer proteins and clearance by active mechanisms, such as extrusion through plasma membrane pumps or ER re-uptake, would lower the free Ca^2+^ in the cytoplasm still further (50). Our basic model does not include these effects and thus sets a conservative bound on cytosolic Ca^2+^ levels.

The Ca^2+^ capacity of the ER lumen is greatly enhanced by the presence of Ca^2+^-binding buffer proteins, such as calreticulin, PDI, GRP94, and others (51). Many of these buffer proteins contain domains with high capacity and low affinity for Ca^2+^ storage. For example, each calreticulin protein contains approximately 25 Ca^2+^ binding sites (47, 52). We take the buffer protein concentration in the lumen to be approximately 0.1mM, based on prior measurements of calreticulin densities (53). The steady-state free [Ca^2+^]_ER_ is set to *U*_0_ *≈* 0.5mM, a typical value within the range of published values (54–56), and in agreement with our own measurements (see below). Thus, the concentration of buffer binding sites is approximately 5*×* greater than the free Ca^2+^ concentration. Because only soluble ions can be released through channels in the ER membrane, the overall dynamics of ER Ca^2+^ release depends on the interplay between buffer-bound and unbound Ca^2+^.

We assume that Ca^2+^ binding is equilibrated at all times – employing a rapid buffering approximation that has been previously validated for modeling Ca^2+^ puff dynamics (57). The evolution of free Ca^2+^ concentration (derived in SI Methods) can then be expressed as:

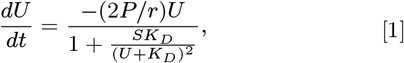

where *U* is the free [Ca^2+^]_ER_, *S* is the total concentration of buffer sites, *K*_*D*_ is the equilibrium dissociation constant, *P* is the effective membrane permeability, and *r* is the ER tubule radius. This expression is integrated to give the free Ca^2+^ levels in the lumen over time, and hence the Ca^2+^ release current *J* (*t*) = 2*πrLPU* (*t*) from the well-mixed local ER region.

To model the local Ca^2+^ concentrations in the cytoplasm, we assume that Ca^2+^ ions diffuse outward from a point-like release source, and that the measured cytoplasmic concentration is averaged over a sphere of radius *R*_loc_ surrounding the source (details in SI Methods). The cytoplasmic Ca^2+^ concentration profile (Fig. 1b) displays a peak at very short times (few ms), followed by a steep decline. This decline arises from a combination of Ca^2+^ depletion in the lumen and the dilution effect of the released Ca^2+^ spreading out over ever-larger volumes. We do not include the effect of active pumps that remove Ca^2+^ from the cytoplasm, which would lower the cytoplasmic levels still further. Notably, plugging in physiologically relevant parameters implies that the maximum released Ca^2+^ concentration is generally sub-*μ*M, orders of magnitude lower than the Ca^2+^ concentration within the ER.

Cytoplasmic Ca^2+^ concentrations drop sharply over subsecond time-scales. After a time of 0.1 seconds post-release, the local Ca^2+^ concentration is expected to be below 20nM regardless of the permeability for releasing Ca^2+^ from the ER. We note that binding to cytoplasmic buffer proteins is known to slow the spreading of Ca^2+^ in the cytoplasm (58, 59). However, such binding must also reduce the free Ca^2+^ levels surrounding the release site (49). In SI Appendix (Fig. S1), we show that cytoplasmic buffering does not substantially raise the local free Ca^2+^ concentration. The minimalist model presented here thus sets a limit on cytoplasmic Ca^2+^ concentrations following a purely local release event.

The magnitudes of localized Ca^2+^ release events from the ER have been estimated to range from *∼* 30nM (for single-channel ‘blips’), to *∼* 200nM (for ‘puffs’ in oocytes and ‘sparks’ in cardiomyocytes) (54, 60–62), to puffs of up to 1*μ*M in osteoclasts (63). Furthermore, the spatial extent of the Ca^2+^ releasing region in puff events has been estimated as diffraction-limited (*R ≈* 0.25*μ*m) (64, 65). Using our estimated parameters, the total amount of luminal Ca^2+^ in a region of this size around an ER junction is only about 10^4^ ions. By contrast, the amount of Ca^2+^ released in oocyte blips and puffs has been estimated at 10^5^ *−*10^6^ ions (65). Achieving these transient elevations of cytoplasmic Ca^2+^ would require a complete loss of the local luminal pool, necessitating transport through the ER.

While we use a modest estimate for the luminal Ca^2+^ carrying capacity, increasing the density of binding sites by an order of magnitude (to *∼* 1mM buffer protein concentration) is still not sufficient to raise cytoplasmic levels above 200nM (SI Appendix, Fig. S2). Thus the locally available pool of Ca^2+^ is insufficient to sustain local release events, and the ions must tunnel through the lumen from neighboring regions of the ER network.

### Network transport and moderate buffer strength combine to enable sustained release

Since local Ca^2+^ is insufficient to produce measured puff magnitudes, we next consider the effect of transport through the lumen of the ER network. We extend our initial well-mixed system into a spatially-resolved reaction-diffusion model for Ca^2+^ ions and buffer proteins within a network of tubules with a locally permeable release region. The peripheral ER in adherent mammalian cells forms a highly looped lattice-like planar network structure (Fig. 1a). We represent this morphology with a honeycomb network (Fig. 1c) with edge length *£* = 1*μ*m (66, 67) and tubule radius *r* = 50nm (12, 68). A local release region, with permeability *P* for free Ca^2+^ ions, is taken to have a radius of *R* = 0.25*μ*m, as suggested for measurements of Ca^2+^ puffs in *Xenopus* oocytes (64).

Both free Ca^2+^ ions and buffer proteins are assumed to diffuse through the ER tubules. The diffusion coefficient of molecular components within the crowded lumen of the ER is not well-established, with prior work indicating that protein diffusivity may be 2 *−* 10-fold lower in the lumen than in the cytoplasm (69–71). We estimate luminal protein diffusivity as *D*_*b*_ *≈* 3*μ*m^2^*/*s, based on our analysis of photoactivated protein spreading (SI Appendix, Fig. S4). Free Ca^2+^ ions are assumed to diffuse an order of magnitude faster (59), with *D*_Ca_ *≈* 30*μ*m^2^*/*s. The reaction-diffusion equations relating concentrations of free ions *U* (*x, t*), total ions *C*(*x, t*), and total buffer sites *S*(*x, t*) are:

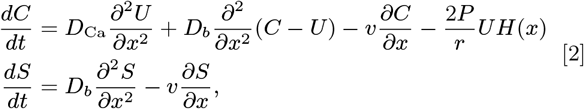

where *H* is an indicator function equal to 1 in the permeable region and 0 otherwise, and the advective term is removed (flow velocity *v* = 0) for purely diffusive transport. For this purely diffusive case, the concentration of total buffer sites *S* is spatially constant at all times. The case with included advective flow within the ER lumen is considered in the subsequent section. We note that the total ion concentration *C* and the unbound ions *U* are not independent quantities, and are related through the rapid-equilibrium approximation (57) according to:

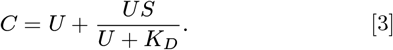

Equations 2 and 3 are solved numerically using a finite volume method that evolves forward the two fields *U* (*x, t*) and *S*(*x, t*), as described in SI Methods. In treating diffusion along the tubules with a single coordinate, we assume that the equilibration of concentration profiles across the cross-section of the narrow ER tubules is very rapid (*<* 1ms equilibration time with the parameters used here).

The release of Ca^2+^ from the network proceeds in several phases (Fig. 1d, SI Video 1). At very short times the local luminal free Ca^2+^ levels are maintained by release from the buffer sites. Next, there is a subsecond time period during which Ca^2+^ is transported through effectively onedimensional (1D) tubules to the permeable zone. Standard solutions of the 1D diffusion equation imply that the concentration near an absorbing region scales as *∼ t*^*−*1*/*2^ (72). On longer time periods of roughly 0.1 *−* 10sec, delivery of Ca^2+^ to the release site involves transport over the two-dimensional (2D) network. This leads to local concentrations that drop as 1*/* log *t*, so that the luminal free Ca^2+^ is very slowly depleted over this range of time scales.

The Ca^2+^ delivery dynamics can also be seen by plotting the total amount of Ca^2+^ released (Fig. 1e), where a rapid release over millisecond timescales is followed by a release rate that changes very slowly over the course of multiple seconds. The cytoplasmic Ca^2+^ surrounding the release site (Fig. 1f) demonstrates a peak at tens of milliseconds, followed by a slow decay as 2D transport of Ca^2+^ through the network does not keep up with 3D dilution in the cytoplasm. Notably, the levels of cytoplasmic Ca^2+^ at second timescales in this model are an order of magnitude higher than those predicted for purely local Ca^2+^ release (Fig. 1b vs 1f), highlighting the importance of transport through the network.

We note that these simulations center the release region on a junction node. In reality, channel opening and release could occur anywhere along the peripheral ER, both close to the junctions and in the middle of longer tubules. For the lattice network with 1*μ*m tubule lengths, we find that the release rate in the middle of a tubule differs from that at a junction by only about 10% after an initial sub-second transient period (SI Appendix, Fig. S5). This observation further highlights the point that transport to the release region limits the Ca^2+^ flux, which is not simply proportional to the luminal volume within the permeable zone.

A key assumption of our model is that the initial concentration of free Ca^2+^ in the ER lumen (*U*_0_) is kept constant regardless of the buffer strength, which in turn modulates the total Ca^2+^ storage capacity. This assumption relies on the notion that the various mechanisms responsible for maintaining luminal Ca^2+^ homeostasis will respond to free Ca^2+^ rather than total Ca^2+^ concentrations. An important consequence is that as the buffer site binding becomes stronger (lower *K*_*D*_), the total initial Ca^2+^ in the system increases, up to the maximum carrying capacity defined by the concentration of buffer binding sites. The amount of Ca^2+^ released over time has a non-trivial dependence on the buffer strength. When the buffer strength is well-matched to the free Ca^2+^ concentration (*K*_*D*_ *≈ U*_0_), the initial levels of free Ca^2+^ can be robustly maintained in the lumen as the release proceeds. However, stronger binding (lower *K*_*D*_) enables a greater total Ca^2+^ storage capacity in the ER so that there is more Ca^2+^ to be released.

Overall, an optimum intermediate value 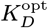 maximizes both the amount of released Ca^2+^ and its cytoplasmic elevation. This optimal value is robust over a broad range of probe times during which the release is measured (Fig. 1e, inset). In the initial phase of rapid release, 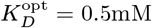 provides optimal buffering for the localized release region. In the infinite-time limit, when the lumen becomes fully depleted of Ca^2+^, an arbitrarily small value of *KD* becomes optimal since it allows storage of the maximum amount of total Ca^2+^ in the lumen. However, in the transport-dominated phase, the local concentration of free Ca^2+^ decreases slowly as the effectively two-dimensional transport through the network replenishes the leakage in the permeable region. Over a range of 0.1 *−* 10sec, an intermediate value of *KD ≈* 0.2mM is optimal both for maximum Ca^2+^ release and maximal cytoplasmic Ca^2+^ elevations (Fig. 1e,f). The effect of the optimal *KD* is enhanced if there is a higher density of luminal buffer binding sites than the modest estimate used here (SI Appendix, Fig. S3). The value of the optimal binding strength is largely independent of the assumed buffer site density. Its dependence on the permeability and size of the local release region is described in SI Appendix Fig. S6.

Notably, the luminal Ca^2+^-binding protein calreticulin contains high-capacity Ca^2+^ binding sites with a moderate binding affinity (*K*_*d*_ *≈* 0.25mM (73)). This value matches well to the optimal binding strength found in our model system. The model thus provides a potential explanation for the limited binding strength of these Ca^2+^ buffers, by demonstrating the increased efficiency of local Ca^2+^ release associated with relatively weak binding.

### Buffer protein mobility enhances Ca^2+^ release

The transport of Ca^2+^ ions through the extended ER lumen to the local release site involves a combination of rapid free ion diffusion, and the slower diffusion of the Ca^2+^-binding buffer proteins which carry the majority of luminal Ca^2+^. Although much slower than the Ca^2+^ ions, the mobility of buffer proteins does contribute to the flux of Ca^2+^ release, on timescales beyond the initial emptying of the local region (Fig. 2a). These results are consistent with the previously noted enhancement of local Ca^2+^ currents by diffusive buffer proteins (38, 74). The effect of buffer diffusivity is found to be particularly important when buffer strength is tuned to its optimal value (Fig. 2b).

**Fig. 2.**
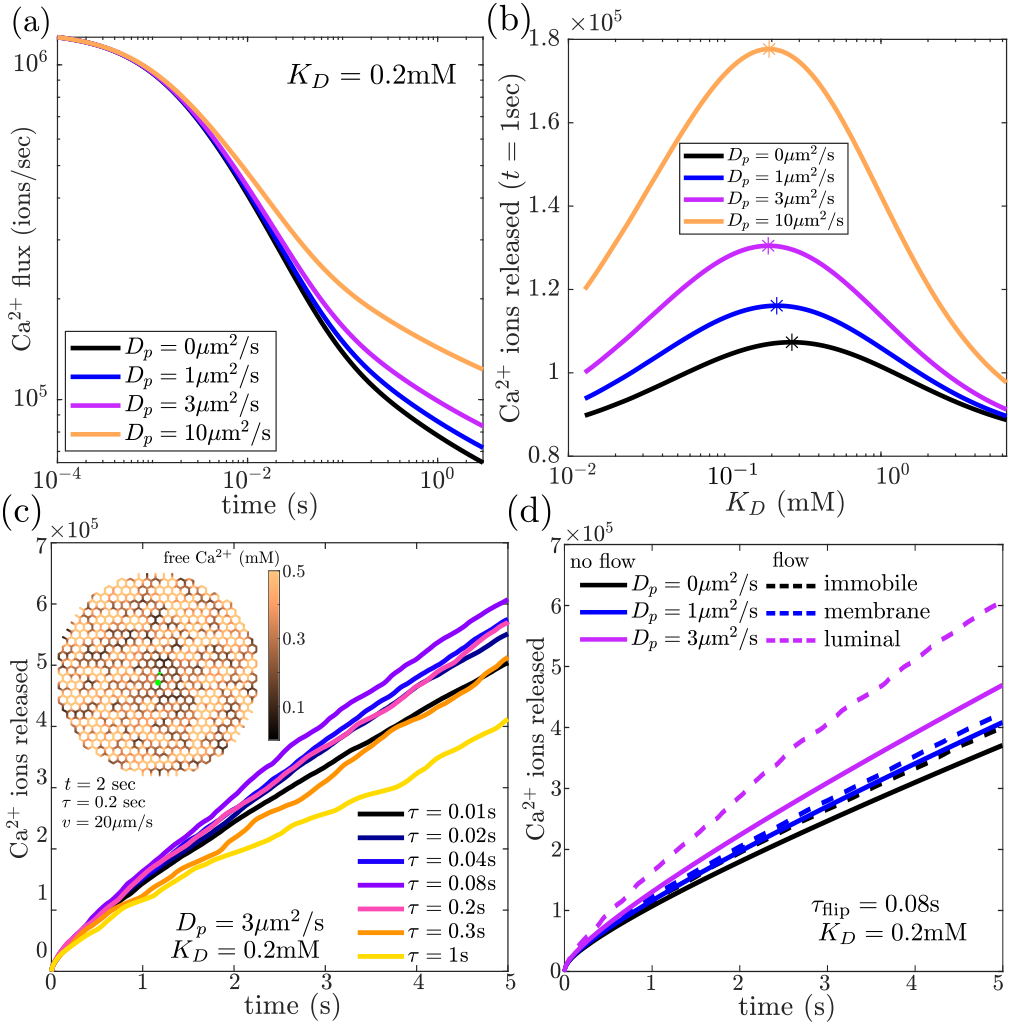
Mobility of buffer proteins and active luminal flow can enhance local Ca^2+^ release in spatially resolved model. (a) Ca^2+^ flux out of local release region in a honeycomb network, for different values of buffer protein diffusivity. (b) Cumulative Ca^2+^ ions released in a 1sec interval, as a function of buffer binding strength *K*_*D*_, for different protein diffusivities. (c) Cumulative Ca^2+^ released over time in an active network with flow velocity *v* = 20*μ*m/s, and different persistence times *τ*_switch_, demonstrating maximum release with flows of intermediate persistence. Each curve is averaged over 10 independent realizations of dynamic random flow patterns. Inset shows simulation snapshot, with green circle marking release region. (d) Cumulative Ca^2+^ released over time for different values of buffer diffusivity, with active flows (dashed lines) or without them (solid lines). Black and blue curves correspond to immobile and membrane-bound buffer proteins, which are not subject to luminal flow.

Recent measurements of single-particle trajectories for luminal proteins have indicated that such proteins can traverse individual ER tubules much more rapidly than would be expected from diffusion alone (75). The observed dynamics of luminal particles is consistent with persistent randomly-directed runs across individual ER tubules, possibly associated with short-range luminal flows, giving rise to a model of the ER as an active network (76). Active motion through the ER lumen is further supported by the apparently superdiffusive spatiotemporal spreading of photoactivated luminal proteins (12) (see also SI Appendix, Fig. S4). Although the mechanism underlying this behavior is not currently well established, we investigate the modulation of Ca^2+^ transport and release by effectively advective motion along the tubules. Specifically, we consider randomly directed flows of velocity *v*, which flip direction independently for each individual edge, as a Poisson process with timescale *τ*. Particles move through this network with a combination of diffusive motion and advection by the edge flows. The rapid flow velocity along the edges is set to *v* = 20*μ*m/s, as previously reported for short-range processive movements of luminal proteins in ER tubules (75).

A dimensionless quantity that describes the importance of flow versus diffusion is the Péclet number: Pe = *vL/D*, where *v* is the flow velocity, *D* the diffusivity, and *L* a length scale of interest. A value of Pe *»* 1 indicates that transport is primarily driven by flow, while Pe *«* 1 implies that diffusion is dominant and the flow makes no noticeable difference over that length scale. For the case of free Ca^2+^ ions moving over a single network edge, we estimate Pe *≈* 0.7, implying that the putative luminal flows will have little effect on the free ion transport. However, for buffer proteins, the corresponding Péclet number is an order of magnitude higher (Pe *≈* 7). Consequently, we would expect advective flow of the luminal contents to substantially enhance the spreading of Ca^2+^-binding buffer proteins but not the free ions themselves.

In Fig. 2c we plot the cumulative release of Ca^2+^ in a network with random advective flows along its tubules (Fig. 2c inset, SI Video 1). We see that flows with a moderate persistence time of *τ ≈* 0.08s allow for the most rapid local Ca^2+^ release. Similar persistence times are shown to optimize the mean squared displacement of particles in an active network (SI Appendix, Fig. S7), by allowing the particles to rapid traverse individual edges without becoming trapped at convergent nodes (76).

By contrast, when buffer proteins are immobilized, the Ca^2+^ release is substantially slower and largely unaffected by flow (Fig. 2d). Similar results are seen if we consider the buffer proteins to be embedded in the ER membrane, which would be expected to give them a lower diffusivity (*D*_*b*_ *≈* 1*μ*m^2^*/*s (77)) and leave their motion unaffected by any putative luminal flows. Thus, luminal flows enhance Ca^2+^ release specifically by driving mobile luminal buffer proteins at high Péclet number, with little effect on the free Ca^2+^ ions themselves.

These results highlight the importance of buffer protein mobility for efficient Ca^2+^ delivery. For luminal proteins, this mobility can be further enhanced by super-diffusive motion in the tubules. Notably, the luminal chaperone calreticulin binds Ca^2+^ with high capacity through the many negatively charged residues of its acidic C-terminal domain (52). However, the closely related membrane protein calnexin has its C-terminal domain situated on the cytosolic side of the ER membrane and thus lacks the ability to provide high-capacity Ca^2+^ storage (78). Our calculations imply that by relegating Ca^2+^ buffering capacity to mobile luminal proteins, the cell may gain an advantage in the ability to rapidly release Ca^2+^ from local regions of the ER.

### ER structural perturbations reduce luminal connectivity and hinder transport

Our physical model demonstrates that longrange transport of Ca^2+^ ions and mobile buffer proteins through the ER lumen is key to efficient localized Ca^2+^ release. These results highlight a potentially important role for the highly connected morphology of the peripheral ER as an intracellular transport network. A natural empirical test of this theoretical prediction would involve perturbing the ER architecture to limit the connectivity of the peripheral network, and measuring the consequent effect on luminal particle transport. However, the genetic and pharmacalogical toolkit for altering ER structure is limited. We use state-of-the-art manipulations relying on the altered expression of two ER tubule morphogens to affect visible changes in the network.

An extreme change in network connectivity is achieved through overexpression of the tubule-stabilizing protein RTN3e, resulting in the peripheral ER lumen breaking up into poorly connected seemingly vesicular structures (79) (Fig. 3a). To quantify the spreading of luminal material through the perturbed ER, we use continuous photoactivation followed by spatial chase (CPAC) (12) of ER-targeted photoactivable GFP (paGFP^ER^) from an initial activation region (Fig. 3b, SI Video 2). The rate of signal rise is recorded within small regions of interest (ROIs) at different distances from the photoactivation site, and the median time-scale for signal arrival is found to increase with increasing distances (Fig. 3c). Notably, the eventual arrival of photoactivated luminal proteins in spatially distant ROIs implies that some luminal connectivity is maintained even in the apparently fragmented ER of RTN3 OE cells. However, the arrival times are much longer in RTN3 OE cells as compared to WT cells (Fig. 3c,d), where we observe a super-diffusive transport profile for paGFP^ER^ luminal marker as has been shown to be consistent with an active flow model of ER luminal transport (12, 75). The greatly reduced arrival rates of luminal proteins may be due to slow escape from voluminous ER fragments into narrow tubular connections, or to constrictions of tubules that serve to partially block luminal continuity between the ER fragments. Such constrictions have previously been reported to arise from overexpression of reticulon proteins (80).

**Fig. 3.**
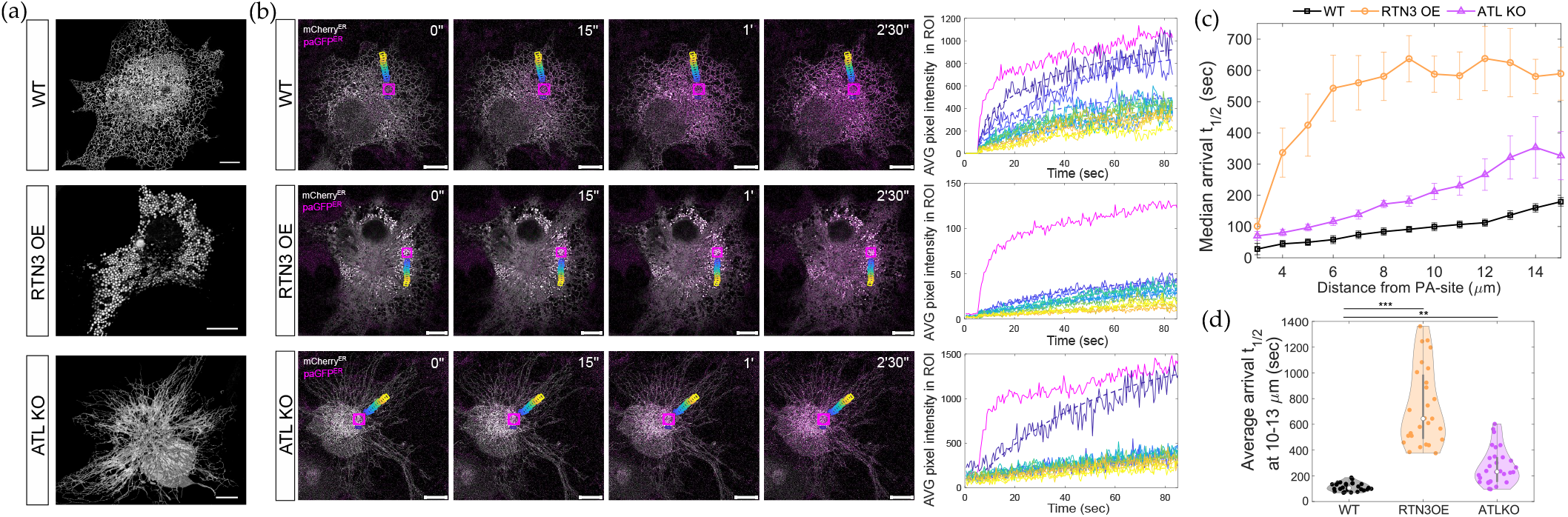
Live-cell imaging shows reduced transport of luminal proteins in perturbed ER morphologies. (a) Representative image of exogenously expressed ER luminal marker (mCherry^ER^) in live COS-7 cells WT and overexpressing an ER shaping protein reticulon 3 isoform E (RTN3 OE) or double knockout of Atlastin 2 & 3 (ATL KO). Note the fragmentation of lumen into enlarged, non-reticular structures and reduction in number of junctions along long, unbranched tubules respectively. (b) Photoactivation pulse-chase assay. Illumination of ER-targeted photoactivable GFP (paGFP^ER^) with 405nm laser restores the characteristic fluorescence of conventional GFP. Spatiotemporal spreading of the photoactivated paGFP^ER^ molecules in the ER lumen is monitored while being continuously and locally supplied at photoactivation site (PA-site) in the perinuclear region during image acquisition. ER luminal transport rates are deduced from quantifying the effective half-time for signal rise of the paGFP^ER^ at region of interests placed at increasing distances around the PA-site. Representative merged images of exogenously expressed ER luminal marker (mCherry^ER^) and paGFP^ER^ signal distribution in COS-7 cells with normal ER network (WT), fragmented lumen (RTN3 OE) and unbranched tubules (ATL KO). PA-site is represented in magenta together with representative ROIs along one axis around it. Corresponding paGFP^ER^ average (AVG) pixel intensity in those ROIs over time is plotted on the right panel. (c) Median paGFP^ER^ arrival time at respective distances from the PA-site (averaged together over all cells) (d) Average arrival time in ROIs located 10-13*μ*m around PA-site. Each dot represents the arrival times from all ROIs at a specific distance in a single cell, averaged together. [WT: n= 8, RTN3 OE: n=8, ATL KO: n=14 cells. ^**^*p <* 0.01, ^***^*p <* 0.001, KS test performed on distributions of averages for ROIs located at 10*μ*m.] Scale bars: 10*μ*m.

A more moderate alteration to ER connectivity involves manipulating the amount of junctions in the ER tubular network. This is achievable through a double knockout of the junction-forming morphogens Atlastin 1&2 (motor neuron disease-associated ATL2/3), leading to peripheral ER structures with long radially oriented tubules and reduced inter-tubule junctions (81, 82) (Fig. 3a). Analysis of CPAC measurements indicates that luminal content transport is slowed in ATL KO cells, resulting in higher median halftimes of signal arrival to distant regions, as compared to WT cells (Fig. 3c,d). This morphological perturbation thus retains the tubular morphology of the peripheral ER while reducing the overall connectivity and hence its efficacy as a transport network.

Given their demonstrated effect on protein transport rates, the manipulations of ER structure by RTN3 overexpression and ATL knockout provide a practical approach for probing the dependence of Ca^2+^ signaling on ER integrity, both *in silico* and in live cells.

### Reduced ER connectivity is predicted to lower local Ca^2+^ release rates

We next proceed to investigate the extent to which altered ER architecture and the concomitantly reduced luminal transport should be expected to alter Ca^2+^ release magnitudes. To represent the concrete changes in ER morphology associated with morphogen perturbations, we extract local 14*μ*m-diameter regions of ER network structure, defined by junction nodes connected via one-dimensional curved edges, from both WT and ATL knockout cells (Fig. 4a.i, a.ii). For the RTN3 OE architectures, we extract the location of ER fragments from similar sized regions and simplify the morphology by treating each of the fragments as a spherical ‘bubble’ and assuming that neighboring bubbles are connected by very narrow tubules (Fig. 4b.iii). Although such tubules are not resolved in the ER images, some connectivity between fragments is evident from the long-range spreading of photoactivated signal shown in Fig. 3. We take the interbubble tubules to have default radius *r*_*b*_ = 10nm, setting them to be substantially narrower than the well-resolved tubules of WT ER.

**Fig. 4.**
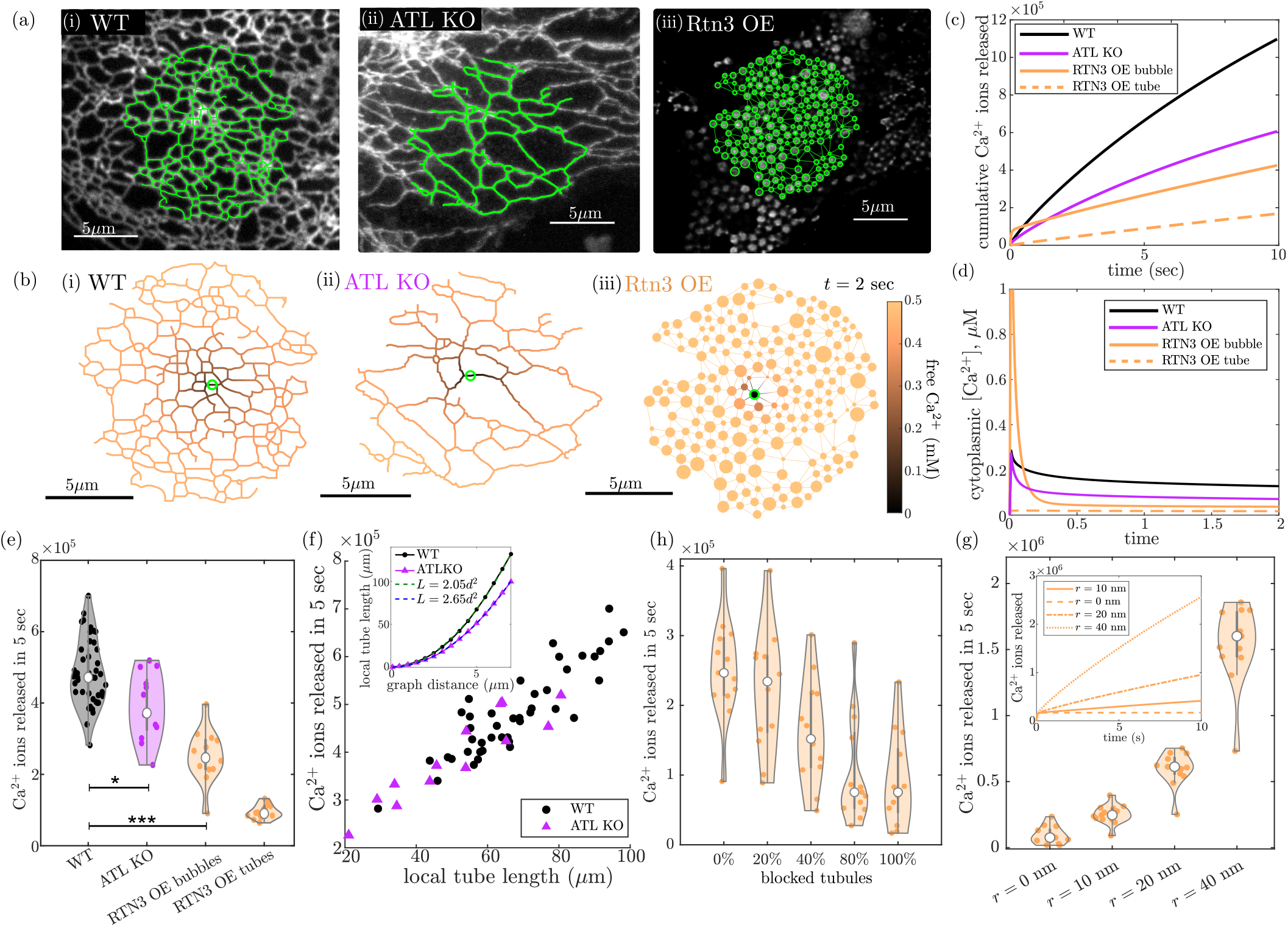
Model of Ca^2+^ release predicts that ER network architecture modulates local release magnitude. (a) Network structures are extracted from sample regions of (i) WT, (ii) RTN3 OE, and (iii), ATL KO cells. Vesicular fragments in RTN3 OE cells are assumed to be connected by 10nm-radius tubules. (b) Simulation snapshots of free Ca^2+^ concentration at *t* = 2sec after initiation of release. Green circle marks release region. (c) Simulated cumulative Ca^2+^ ions released over time from the sample network structures shown in (b). For RTN3 OE structure, solid line gives result for release centered on a bubble region and dashed line has release centered on tubule. (d) Predicted cytoplasmic [Ca^2+^], averaged over 1*μ*m region around release site, for same simulation runs as in (b). (e) Cumulative Ca^2+^ released after 5 sec, for distinct 14*μ*m-diameter regions of WT (*n* = 44, from 22 cells), RTN3 OE and ATL KO (*n* = 13, from 4 cells) networks. (*) *p* = 0.015; (***) *p <* 10^*−*4^, by two-sample Kolmogorov-Smirnov test. (f) Cumulative Ca^2+^ release plotted against the local tube length, within 5*μ*m graph distance of the release center, for WT and ATL KO network regions. Inset: Local tube length within graph distance *d*, plotted as a function of *d*, for WT and ATL KO network regions. Dashed lines show fit to quadratic scaling as expected for effectively 2D networks. (g-h) Effect of reduced connectivity on cumulative Ca^2+^ ions released over 5sec from RTN3 OE network structures as in (e). (g) Different tubule radii *r*. Inset shows ions released over time. (h) Different percentage of tubules blocked. Parameters are set to *D*_*b*_ = 3*μ*m^2^*/*s, *K*_*D*_ = 0.2mM, *P* = 20*μ*m/s, *R* = 0.25*μ*m throughout.

For each network structure, a point along the network edges is selected close to the center of the region to serve as the locus for localized Ca^2+^ release. For the partially fragmented RTN3 OE networks, we consider both release from a bubble and from the middle of a connecting tubule. As before, the size of the permeable region is taken to be *R* = 0.25*μ*m. This approximation rests on prior published work demonstrating that Ca^2+^ puffs originate from diffraction-limited regions (64, 65). Larger release regions are considered in SI Appendix (Fig. S8) and shown to not qualitatively alter the results.

For simplicity, we assume that the initial concentration of free Ca^2+^ and buffer proteins is the same in all of the network structures (substantiated by measurements described below). Furthermore, we take the same value of the permeability *P* = 20*μ*m/s throughout. Taken together, these assumptions enable us to directly compare the role of altered network morphology in modulating Ca^2+^ release, while keeping other parameters constant.

Using our numerical simulations of luminal Ca^2+^ dynamics, we compute the release of Ca^2+^ ions over time from the permeable regions in the different network architectures. The bubbled fragments in the RTN3 OE networks are treated as well-mixed reservoirs, with narrow-escape kinetics governing the release of Ca^2+^ ions into the adjacent tubules (details in SI Methods). As can be seen in the simulation snapshots (Fig. 4b, SI Video S3), the local depletion of Ca^2+^ extends to a larger spatial distance in the ATL KO networks, with their long tubules and sparser junctions, as compared to the denser WT networks. This leads to slower Ca^2+^ release from the mutant networks (Fig. 4c). For the RTN3 OE morphology, the releasing bubble becomes rapidly depleted, so that at very short (millisecond) timescales, the predicted Ca^2+^ release is larger in these networks. However, the delivery of ions and buffer proteins through the very narrow connecting tubules is quite slow and the overall Ca^2+^ release over seconds timescales is lower than in WT. Similar trends are observed for the predicted cytoplasmic Ca^2+^ levels (Fig. 4d). If the release in the RTN3 OE network occurs from the middle of a tubule rather than a bubbled fragment, the Ca^2+^ flux is predicted to be much lower than for WT networks on all timescales.

Figure 4e demonstrates that the total amount of Ca^2+^ released in 5 sec is significantly lower in ATL KO networks and partially fragmented RTN3 OE structures, as compared to WT, when compared across multiple extracted architectures. We note that the broad spread in different release magnitudes is due entirely to the varying architectures of individual network regions within each group of cells. Comparison of individual ATL KO and WT network regions demonstrates that the amount of Ca^2+^ released correlates with the local tube length (or equivalently the luminal volume) available within a short graph distance of the release point (Fig. 4f). These results are consistent with prior work demonstrating that structural heterogeneity of the ER can play an important role in modulating transport through the network (83).

The available local edge length scales quadratically with increasing graph distance for both WT and ATL KO networks, indicating that these networks behave as essentially twodimensional architectures, albeit with reduced density for the ATL KO morphologies (Fig. 4f, inset). The lower release magnitude from ATL KO networks can thus be attributed to the need for bringing Ca^2+^ from greater distances through the sparse network. In SI Appendix (Fig. S9), we show for comparison the simulated release of Ca^2+^ from highly idealized regular lattice morphologies that exemplify the stereotypical structural changes in the perturbed ER architectures. These idealized structures show qualitatively the same effects on Ca^2+^ release as discussed above.

For the partially fragmented RTN3 OE structures, there is a substantially higher volume of ER lumen within the enlarged bubbles, as compared to the WT peripheral ER network. The predicted reduction in Ca^2+^ release occurs despite the increased load of spatially proximal Ca^2+^ ions, as a result of the reduced connectivity through narrow tubules between the bubbles. While we make the conservative assumption that buffer protein concentration remains constant in the perturbed networks, it is plausible that the total amount of buffer protein manufactured by the cell would remain constant instead. In that case, the increased ER luminal volume would correspond to a diluted buffer protein concentration, with a concomitant slight lowering of the Ca^2+^ release magnitude (see SI Appendix, Fig. S10).

The Ca^2+^ delivery rates in the fragmented RTN3 OE structures can be tuned by adjusting the connectivity between neighboring bubbled fragments. For example, wider tubules connecting the bubbles allow for substantially faster Ca^2+^ release (Fig. 4g). Extremely narrow tubules resulting from RTN3e overexpression may also be subject to sporadic blockages, where closure of the tubule prevents luminal connectivity (80). As the extent of tubule blockage is unknown, we explored a range of possible blockage probabilities (Fig. 4h), which showed the decrease in Ca^2+^ release as more tubules are blocked. Completely disconnecting the bubbles results in all of the Ca^2+^ being released from the permeable bubble within approximately 0.1sec (Fig. 4g, inset), followed by no further release. Overall, the simulation results on these highly perturbed fragmented networks highlight the importance of Ca^2+^ tunneling through a well-connected ER lumen for sustaining a high magnitude of release.

To further explore the influence of ER structure on its Ca^2+^ release capacity we extended our model beyond purely tubular ER to account for ER sheets, which constitute one of the two main interconnected domains of the organelle. ER sheets may serve as enlarged reservoirs of luminal volume to support nearby Ca^2+^ release. Analyzing sheet-rich regions extracted from WT cells, we observe that Ca^2+^ releases in the vicinity of sheets are slightly higher in magnitude, due to the greater amount of Ca^2+^ available near the release site (Fig. S11). However, the increase is relatively modest, highlighting the importance of limited transport through narrow tubules and not just the nearby luminal volume in determining Ca^2+^ release magnitude.

### Ca^2+^ release is slowed in perturbed ER morphologies

We next proceed to experimentally quantify local Ca^2+^ release events in COS-7 cells with functional ER Ca^2+^ storage but varying extent of network connectivity, with either no perturbation (WT), highly disconnected networks in RTN3 OE cells, or mildly reduced network connectivity in ATL KO cells.

First, we establish the extent to which ER morphological manipulations affect the steady-state ER Ca^2+^ concentration and its overall storage capacity (parameters that can influence the magnitude of ER release). To measure ER Ca^2+^ in absolute terms we used a FRET reporter with a relatively low *K*_*d*_ tuned to the luminal high Ca^2+^ environment (84), read out by fluorescence lifetime imaging (FLIM). We improved its optical properties, extending the accuracy in calibrated FLIM-based measurements, by optimizing the photophysical characteristics of the FRET donor/acceptor (mTurquoise with a single lifetime and REACH with low emission, Fig. 5a; details in SI Methods). The dynamic range of the new probe is well adapted to detect physiological variations in ER Ca^2+^ concentration ([Ca^2+^]_ER_) after calibration in maximal and minimal Ca^2+^ conditions. Measurements in COS-7 WT confirm the baseline concentration [Ca^2+^]_ER_ = 0.47*±*0.11 mM (Fig. 5b, c). The structurally perturbed ER retained its Ca^2+^ storage levels: free Ca^2+^ concentrations were 0.59*±*0.19 mM and 0.62*±*0.16 mM in cells with tubular junction loss (COS-7 ATL KO) and in the fragmented ER (COS-7 RTN3 OE), respectively. Despite the large-scale perturbations to ER structure, it thus retains its ability to function as a Ca^2+^ storage reservoir, without substantial changes in the overall concentration of luminal free Ca^2+^.

**Fig. 5.**
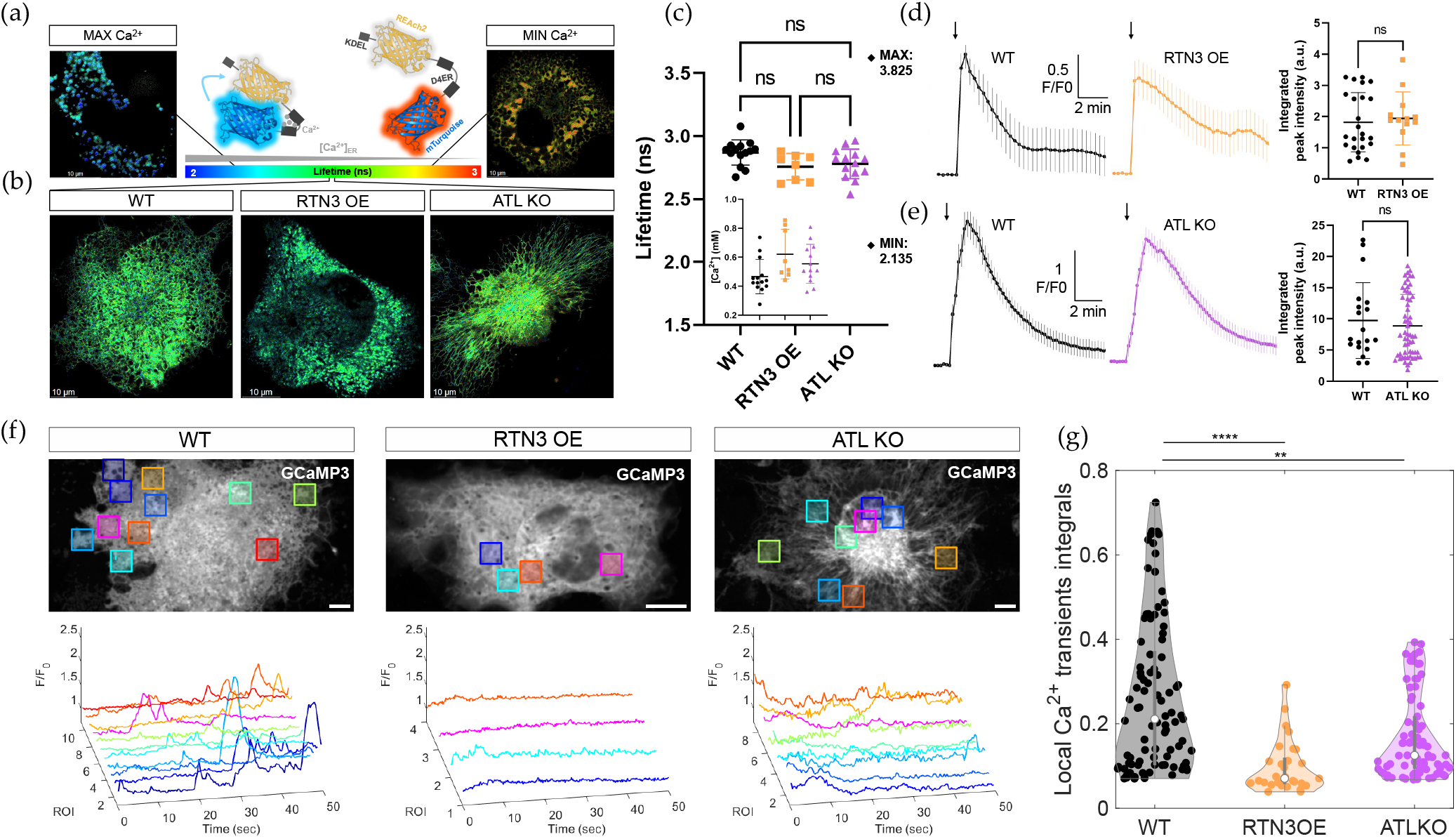
Perturbation of ER architecture hinders localized Ca^2+^ transients without affecting Ca^2+^ storage in live cells. (a) (center) Schematic of FLIM method for measuring absolute Ca^2+^ concentration; ER-targeted D4ER-Tq FRET Ca^2+^ probe with new fluorophores is drawn bound/unbound to Ca^2+^ (high/low FRET respectively) with corresponding mTurquoise’s lifetimes. KDEL: ER retention motif. Representative FLIM images for calibration, showing extreme luminal Ca^2+^ levels with (left) maximal Ca^2+^ obtained by ionomycin (10*μ*M) treatment and (right) minimal Ca^2+^ obtained by thapsigargin (3*μ*M) treatment. (b) Representative FLIM images in COS-7 cells with altered ER network structure corresponding to WT, RTN3 OE and ATL KO. (c) Quantitative FLIM measurements of luminal Ca^2+^ concentration (each data point represents one cell). Mean [Ca^2+^]_ER_ is not significantly changed between conditions. Mean lifetimes in maximal and minimal Ca^2+^ conditions are indicated. (WT: n=14, RTN3 OE: n=8, ATL KO: n=14; ns: *p >* 0.05, 1-way ANOVA). (d-e) Measurement of total ER Ca^2+^ load by global depletion. Cytosolic Ca^2+^ fluorescence intensity is plotted over time after addition of thapsigargin (1*μ*M, indicated by the arrow). Integrated fluorescence intensity is not significantly changed between conditions. Probes used were (d) Oregon Green BAPTA-1, AM for WT/RTN3 OE (WT: n=24, RTN3 OE: n=13, ns: *p >* 0.05, KS test) and (e) GCaMP3 for WT/ATL KO (WT: n=19, ATL KO: n=59. ns: *p >* 0.05, KS test). (f) Representative fluorescence intensity images of GCaMP3 probe for cytoplasmic Ca^2+^ released from the ER in COS-7 WT, RTN3 OE and ATL KO cells after light-induced IP_3_ uncaging. Semi-automatically detected ROIs around local Ca^2+^ transients are represented as colored squares and corresponding signal intensity traces over time within each ROI are plotted. (g) Quantification of integrated fluorescence intensity under each peak during local Ca^2+^ transients (each data point represents one local Ca^2+^ transient from WT: n=18, RTN3 OE: n=10, ATL KO: n=25 cells. ^****^*p <* 0.0001,^**^*p <* 0.01, KS test).

Further, the total ER Ca^2+^ load appears not significantly altered in cells with fragmented (RTN3 OE) or junction-depleted (ATL KO) ER architecture compared to COS-7 WT, as evident from quantifying the total release elicited by thapsigargin (a SERCA pump inhibitor that globally releases free and buffer-bound ER Ca^2+^; Fig. 5d, e). Thus, the membrane components regulating Ca^2+^ content in the ER (Ca^2+^ pumps and channels (85)) appear to not be significantly impaired by ER morphological changes and effectively prevent the depletion or overload of its Ca^2+^ storage. The live-cell measurements therefore justify our model assumptions that ER morphological perturbation does not reduce either the free or the total Ca^2+^ load in the ER lumen.

An additional factor that could potentially modify Ca^2+^ release rate is the density and clustering of Ca^2+^-releasing channels (IP_3_Rs). The retained ability of cells with perturbed ER to trigger IP_3_-induced Ca^2+^ waves suggests that the channels’ functionality is preserved. To directly assess the status of the Ca^2+^ channels we sought to visualize their ER location and composition. Studying IP_3_R distribution in live cells down to single molecule resolution was recently enabled by developing a cell system with fluorescent tagging of the channels’ endogenous locus (86)). In this system, the spatial distribution of IP_3_R clusters was largely unchanged by the ER morphological perturbations (Fig. S13). Furthermore, as revealed by step-wise bleaching, the typical number of individual channel molecules per cluster was as previously reported ((86)) and remains unchanged in RTN3 OE cells (Fig. S14).

The above measurements demonstrate that networks with perturbed connectivity retain similar baseline levels of Ca^2+^ and similar channel distributions compared to the normally shaped network. We next evaluate the ability of different ER architectures to supply Ca^2+^ in spatially restricted regions. To generate localized Ca^2+^ releases, we use photoactivated IP_3_ uncaging to deliver controlled amounts of IP_3_ by modulating exposure time and intensity of a 405nm laser (details in SI Methods). This triggers sporadic opening of IP_3_R Ca^2+^ channels and generation of localized Ca^2+^ release events (SI Video 4). Ca^2+^ release events are monitored in the cytoplasm by measuring intensity changes of an ER membrane-tethered cytosolic Ca^2+^ sensor (GCaMP3). ROIs with individual release events restricted in space and time are detected semi-automatically (Fig. 5f, details in SI Methods). Each release event corresponds to a defined peak in the local Ca^2+^ signal. This peak is expected to be limited by a combination of several factors: the Ca^2+^ release rate from the ER, the dilution of Ca^2+^ in the cytoplasm, and the chelation and clearance of cytoplasmic Ca^2+^. We thus do not aim to quantitatively compare peak magnitudes with model predictions, but rather to qualitatively note the difference in peaks between cells with differing ER structures.

We estimate the overall extent of release by taking the area under the curve of each transient Ca^2+^ peak. On average, the peak integrals are found to be significantly lower in RTN3 OE cells, suggesting that their poorly connected ER structures reduced the ability to release high amounts of Ca^2+^ in localized regions along the ER (Fig. 5g). Similarly, the peak integrals are reduced for ATL KO cells, where the ER exhibits long tubules and reduced junctions. Although Ca^2+^ release rates cannot be extracted directly from the peak integrals, the reduction in total signal during each transient event is consistent with a decrease in Ca^2+^ release rates.

In addition to the perturbed ER structures considered here, we measured Ca^2+^ release in cells with expanded ER sheets, a phenotype attained through overexpressing Climp63, an ER morphogen responsible for stabilizing sheet structures (87) (Fig. S12a). The cell’s ability to store Ca^2+^ and support local Ca^2+^ releases was preserved upon this manipulation (Fig. S12b-e). ER sheet expansion led to a slight increase in Ca^2+^ release magnitudes (Fig. S12f), as predicted by our model of sheet-like structures interspersed with tubules (Fig. S11).

Overall, the Ca^2+^ release measurements are in line with our model prediction that dense ER tubule connectivity is necessary to support efficient local Ca^2+^ release through transport of Ca^2+^ and buffer proteins from neighboring regions of the network. Perturbations to this connectivity through altered expression of ER morphogens is shown to alter the magnitude of local Ca^2+^ release events. The theory and experiments thus both provide insight into a potentially important functional role of the highly connected ER network morphology.

## Discussion

The physics-based modeling of this study demonstrates that localized Ca^2+^ events rely on Ca^2+^ transport through a well-connected ER lumen, with the aid of mobile buffer proteins. Furthermore, live-cell imaging together with manipulations of ER morphogen expression show that alterations to ER structure both hinder transport through the network and decrease the magnitude of local Ca^2+^ signals.

Raising the local cytoplasmic Ca^2+^ level requires first a one-dimensional ER tubule, then an effectively twodimensional network, to supply sufficient Ca^2+^ to overcome rapid three-dimensional dilution. This functional necessity is achieved by storing a high luminal Ca^2+^ load, primarily bound to high-capacity buffer proteins. Our simulations show that transport through the network results in a very slow depletion of free luminal Ca^2+^ in the release region, giving rise to an optimum intermediate binding strength (*K*_*D*_ *≈* 0.2mM) for the most rapid Ca^2+^ release over a broad range of timescales (0.1 *−* 10 sec). This moderate binding strength matches some estimates of the high-capacity sites of calreticulin, a major Ca^2+^-storing luminal buffer protein (73).

In addition, our model demonstrates that Ca^2+^ release is enhanced by the mobility of the buffer proteins, particularly in the presence of putative active flows within the ER lumen. The nature and mechanism generating such flows is not yet well-established, and may include rapid contraction of ER elements (75, 88) or even chemiosmotic flows coupled to ion dynamics within the lumen. Regardless of the underlying cause, some amount of effectively superdiffusive transport in the ER lumen has been demonstrated via analysis of both single-particle data (75) and photoactivated protein spreading (SI Appendix, Fig. S4). The importance of rapid buffer mobility may explain why the luminal chaperone calreticulin is responsible for carrying much of the intra-ER Ca^2+^ load on its many Ca^2+^-binding sites, while its membrane-bound analogue calnexin lacks a high-capacity Ca^2+^-storage domain (78).

Importantly, our results highlight a key structure–function relationship of the peripheral ER network. The interconnected lattice of ER tubules forms a topologically sequestered domain filled with excess Ca^2+^ in proximity to all cellular regions. Efficient transport through the network allows Ca^2+^ to be rapidly delivered and released anywhere within the cell. We perturb the ER architecture by altering morphogen expression levels, yielding structures with extended tubules (ATL KO) and partially fragmented ER (RTN3 OE). Measurements of photoactivated protein spreading demonstrate that both these structural perturbations result in slower longrange transport through the ER lumen. Our physical model of Ca^2+^ transport predicts decreased rates of Ca^2+^ release in both these altered morphologies. Furthermore, measurements of local Ca^2+^ events magnitudes show that indeed releases are much smaller in the cells with long-tubule or fragmented ER as compared to the WT cells, with their well-connected lattice-like network architecture.

While it is possible that other effects of the ER morphogen perturbations are responsible for the reduced Ca^2+^ release, our additional measurements help eliminate several alternate explanations. In particular, the fluorescence lifetime measurements show that free Ca^2+^ concentration in the perturbed ER lumen is similar to or even higher than the WT cells. Additionally, measuring cytoplasmic Ca^2+^ levels following global release indicates that the total luminal Ca^2+^ load is similar in ATL KO and RTN3 OE cells as compared to WT. Thus, there is no decrease in luminal Ca^2+^ availability to account for the reduced magnitudes of localized Ca^2+^ events. Further, IP_3_R channel distribution and size were not substantially altered by ER structural perturbation. These measurements rule out the possibility that depletion of Ca^2+^ channels on the ER accounts for the substantial decrease in Ca^2+^ release magnitudes. Given that two very different morphological changes to the ER both have in common a demonstrated reduction in ER luminal transport, the most parsimonious explanation is that this reduced transport is responsible for the decreased magnitude of local Ca^2+^ releases, consistent with our model predictions.

In this study, cultured COS-7 cells were employed due to their easily visualized extended ER network, and their tractability as a model system for ER morphogen perturbations. However, the key conceptual results of intra-luminal Ca^2+^ and buffer transport supporting localized release also apply to more specialized cell types. Neuronal cells, in particular, exhibit highly extended ER architectures that provide a continuous Ca^2+^ store linking the cell body to the distal tips of long and/or branched projections (34). In photoreceptor cells luminal transport from the somatic ER has been shown to replenish Ca^2+^ stores in distal termini (89). Perturbations of the ER morphogen Rtn4/Nogo in neuronal cells have been found to modulate axonal regeneration (90, 91) as well as alter Ca^2+^ release (12). Our results demonstrate the need to consider both buffer protein and free ion transport through the extended ER structures of neurons, as well as the importance of robust luminal connectivity. Expansion of the model and live-cell measurements of intra-ER transport to a variety of specialized ER architectures of relevance to neuronal cells forms a potentially fruitful avenue for future work.

In many cellular contexts, localized Ca^2+^ release events (puffs, sparks, or twinkles) play an important biological role in providing the initial trigger for the large-scale signaling waves propagated by Ca^2+^-induced Ca^2+^ release (9). In some cells, frequent localized release events are themselves responsible for functional regulation, as in gliotransmitter release by astrocytes (92). The connection between ER morphology, luminal transport, and Ca^2+^ delivery thus forms a globally important structure-function link at the cellular scale. The intertwined physical modeling and live-cell imaging approaches demonstrated in this work have potentially broad applicability for elucidating the importance of ER architecture in a variety of cellular systems.

## Materials and Methods

### Model for Ca^2+^ buffering and transport

Release and transport of Ca^2+^ ions, in a regime of rapid buffer binding equilibration, is modeled using a finite volume method on networks of narrow tubules, with network structures extracted from images of peripheral ER in COS-7 cells, as described in SI Appendix.

### Cell culture, transfections, constructs, and reagents

COS-7 and EGFP-IP_3_R1 HeLa cells were cultured and transfected, as described in SI Appendix.

### Continuous photoactivation chase (CPAC)

The spatiotemporal dynamics of locally photoactivated paGFP^ER^ was tracked over time in ROIs at different distances from the photoactivation zone, as described in SI Appendix.

### Fluorescence Lifetime Imaging Microscopy (FLIM) and measurement of total Ca^2+^ load

An improved FLIM probe was developed and implemented to measure absolute Ca^2+^ concentrations in the ER lumen. Thapsigargin treatment followed by imaging with GCaMP3, an ER membrane-tethered Ca^2+^ sensor on the cytosolic side, was employed to measure total Ca^2+^ load in the ER. Details of both methods are described in SI Appendix.

### Local Ca^2+^ release event generation and analysis

Local Ca^2+^ release events were generated by photo-uncaging low amounts of caged-IP_3_R3, followed by imaging of cytosolic Ca^2+^ with the GCaMP3 probe and quantitative analysis of spatially localized peaks, as described in SI Appendix.

### Fluorescence microscopy and single-step photobleaching of IP_3_Rs

The spatial distribution of IP_3_R clusters was analyzed via 3D fluorescence microscopy, and cluster size was quantified via single-step photobleaching analysis, as described in SI Appendix.

## ACKNOWLEDGMENTS

We thank members of Colin Taylor’s lab (Cambridge, UK) for kindly gifting the EGFP-IP_3_R1 Hela cell line. EFK is funded by NSF award #2034482, a Cottrell Scholar Award from the Research Corporation for Science Advancement, and a UCSD Chancellor Funded Research Grant. EA is supported by the UK Dementia Research Institute [award number UK DRI-2004] which receives its funding from UK DRI Ltd, funded by the UK Medical Research Council, Alzheimer’s Society and Alzheimer’s Research UK. LMW is supported by NSF award #2034486.

