## Supplemental Material for "Luminal transport through intact endoplasmic reticulum limits the magnitude of localized Ca^2+^ signals"

### SUPPLEMENTARY MATERIAL:
### Luminal transport rates through intact endoplasmic reticulum limit the magnitude of localized $Ca^{2+}$ signals

Cecile C Crapart, Zubenelgenubi C. Scott, Tasuku Konno, Aman Sharma, Pierre Parutto, David M. D. Bailey, Laura M. Westrate, Edward Avezov,* and Elena F. Koslover*

#### S1. SUPPLEMENTAL MATERIALS AND METHODS

##### Model for $Ca^{2+}$ buffering and release

In modeling $Ca^{2+}$ release, we consider both free and buffer-associated $Ca^{2+}$ within the ER lumen. The on- and off-rates of ions from the buffer binding sites are assumed to be very fast, so that the free and bound $Ca^{2+}$ concentrations are always taken to be equilibrated.

This rapid buffering approximation has been previously applied to modeling $Ca^{2+}$ puff dynamics [1]. The approximation is valid so long as the on and off rates are much faster than the time required for $Ca^{2+}$ ions to diffuse over the characteristic length scale of the $Ca^{2+}$ gradient. At the beginning of the simulation, this length scale can be approximated as $\hat{\ell} \approx D_{\text{Ca}}/P \approx 1\mu\text{m}$, with the gradient flattening further over time. The relevant time-scale for rapid buffer equilibration is then $\hat{\ell}^2/D \approx 0.08\text{s}$. Past estimates of buffer binding off-rates are on the order of $100\text{s}^{-1}$ [2, 3], validating the rapid-binding approximation.

Defining $U$ as the free $Ca^{2+}$ concentration, $B$ as the bound $Ca^{2+}$ concentration, $S$ as the total concentration of binding sites, and $K_D$ as the dissociation constant for each binding site, gives the equilibrium relation

$$K_D = \frac{U(S-B)}{B}. \tag{S1}$$

The total $Ca^{2+}$ concentration $C = U + B$ is then

$$C = U + \frac{US}{U + K_D}. \tag{S2}$$

$Ca^{2+}$ release is modeled as occuring in a localized region, containing a total length $L$ of ER tubules with an effective membrane permeability $P$ setting the rate of release for free $Ca^{2+}$ ions. The parameters for our models are summarized in Table S1.

###### *Release of local $Ca^{2+}$ pool in model without transport*

The simplest model is one where the permeable region is completely independent of the rest of the network, and only $Ca^{2+}$ ions within that region can be released. For simplicity, we make the assumption that the permeable region is well-mixed, with no transport processes involved except for the movement of $Ca^{2+}$ ions across the membrane. In this case, we can write the rate of change in luminal $Ca^{2+}$ as

$$\frac{dC}{dt} = -\frac{PA_{\text{ER}}}{V_{\text{ER}}}U = -\frac{2P}{r}U, \tag{S3}$$

TABLE S1. Default parameters for model of luminal $Ca^{2+}$ dynamics

| Parameter | Description | Value | Reference |
|---|---|---|---|
| $U_0$ | initial free [$Ca^{2+}$ ] | 0.5 mM | [4–6] |
| $S$ | concentration of luminal buffer binding sites | 2.7 mM | [7] (500$\mu$g/g cell vol) |
| $D_b$ | buffer protein luminal diffusivity | 2.8$\mu$m$^2$/s | [8] & Fig. S4 |
| $D_{\text{Ca}}$ | free $Ca^{2+}$ luminal diffusivity | 28$\mu$m$^2$/s | set to $10 \times D_b$ [9] |
| $D_{\text{cyto}}$ | cytoplasmic $Ca^{2+}$ diffusivity | 200$\mu$m$^2$/s | [10] |
| $r$ | ER tubule radius | 50$\mu$m | [11] |
| $R$ | spatial extent of local release region | 0.25$\mu$m | [12] |
| $P$ | permeability of local release region | 20$\mu$m/s | [12] (0.4pA peak current) |
| $R_{\text{loc}}$ | radius of region for averaging cytoplasmic $Ca^{2+}$ | 1$\mu$m | |
| $v$ | flow velocity | 20$\mu$m/s | [13] |

where $r$ is the radius of ER tubules, $A_{\text{ER}} = 2\pi rL$ is the surface area of the permeable tubules, and $V_{\text{ER}} = \pi r^2 L$ is their volume. Plugging in Eq. S2 gives the time-evolution of the free $Ca^{2+}$ concentration, as in Eq. 1equation.0.1. The time evolution of luminal free $Ca^{2+}$ is found by integrating Eq. 1equation.0.1. The current of $Ca^{2+}$ out of the ER lumen is then given by

$$J(t) = PA_{\text{ER}}U(t). \tag{S4}$$

To estimate the permeability of the ER within the release region (total tubule length $L = 3R$), we note that measurements of $Ca^{2+}$ puffs in oocytes indicate a typical peak current of approximately $I \approx 0.4$pA [12]. Assuming only local $Ca^{2+}$ is released at very short times, Eq. S4 gives an estimate of the corresponding permeability as $P = J/(2\pi rLU_0) \approx 20\mu$m/s.

To model the local cytoplasmic $Ca^{2+}$ levels, we approximate all the $Ca^{2+}$ release as occuring at a single-point, assume the $Ca^{2+}$ ions spread from this source with diffusivity $D_{\text{cyto}}$, and average the resulting concentration profile over a sphere of radius $R_{\text{loc}} = 1\mu$m. This averaged local cytoplasmic concentration is given by:

$$\psi(t) = \frac{3}{R_{\text{loc}}^3} \int_0^t dt' \int_0^{R_{\text{loc}}} dr\ r^2 G(r, t - t') J(t) \tag{S5}$$

where $G(r, t) = (4\pi D_{\text{cyto}} t)^{-3/2} e^{-r^2/(4D_{\text{cyto}} t)}$ is the Green's function for three-dimensional diffusion.

In addition to this simple estimate of cytoplasmic free $Ca^{2+}$, in the Supplemental Material (Fig. S1) we numerically solve a model that takes into account equilibrated binding of $Ca^{2+}$ ions to diffusive cytoplasmic buffer proteins.

#### Network structures

We obtained WT ER network structures from confocal images of COS-7 cells transfected with 0.2$\mu$g mcherry_KDEL. Cellular transfections, imaging and image processing was carried out as described in prior work [14]. The segmentation toolkit ilastik [15] was employed to segment the ER from the mCherry_KDEL channel, with custom-written skeleton-tracing code (available at https://github.com/lenafabr/networktools) used to extract the network structure. The perinuclear region was manually excised from each image and unphysical terminal nodes arising from segmentation artifacts were manually removed. A total of 44 circular regions of radius $7\mu$m was excised from images of 22 different cells.

For ATL KO cells, the same network extraction procedure was employed as for WT, yielding 13 distinct circular regions from 4 cells.

For RTN3 OE cells, the ER structure was again segmented using ilastik. Individual fragments were fit to circular shapes (using the centroid and average boundary radius around it). A Delaunay triangulation was used to connect neighboring circular fragments, followed by extraction of large circular regions of the network. This procedure yielded 14 non-overlapping network region structures, from 4 different cells. For the simulations, each small circular fragment in the network was treated as a well-mixed spherical reservoir. The edges connecting neighboring reservoirs were treated as tubules of radius 10nm.

For WT and ATL KO network structures, the center of the permeable region was selected by picking the point along the edges closest to the center of mass of the network. In the case of the RTN3 OE networks, the bubble fragment closest to the center of mass was chosen as the release locus. When considering release from tubules, a nearby point half-way along a tubule of length $> 0.5\mu$m was selected as the release region center.

#### Numerical implementation of spatiotemporal dynamics on a network

To incorporate the transport of $Ca^{2+}$ through the ER, we use a numerical finite volume method [16] to evolve forward in time the concentration of free $Ca^{2+}$ ions and buffer sites on a network. Binding of $Ca^{2+}$ to the buffer sites is assumed to be equilibrated in each mesh cell. The finite volume simulations are implemented in Fortran 90 and the source code is provided at https://github.com/lenafabr/networkSpreadFVM.

##### *Finite volume method on tubular network*

The network structure is meshed into cells of length $\Delta l$ (maximum length $0.1\mu$m). Each cell either represents a segment of network tubule or a triskelion of tubules centered around a three-way junction. The network topology defines the boundaries between mesh cells. All mesh cells whose center falls within radius $R = 0.25\mu$m of the network center are labeled as permeable. The surface area of each mesh cell is set to $A_i = 2\pi r\Delta l$ and its volume to $V_i = \pi r^2\Delta l$, where $r$ is the tubule radius.

The state of the network is defined by two field variables: $U_i(t)$ represents the concentration of free $Ca^{2+}$ in mesh cell $i$, and $S_i(t)$ represents the total concentration of buffer sites in mesh cell $i$. Initial conditions are set to spatially constant concentrations $S_i(0) = 2.7$mM and $U_i(0) = U_0 = 0.5$mM. We note that in the case of purely diffusive transport, $S_i(t)$ remains constant for all mesh cells, but in the presence of flows it must also be evolved in accordance with the dynamic equations (Eq. 2equation.0.2). The total $Ca^{2+}$ concentration in a mesh cell ($C_i(t)$) is set at each step using Eq. S2.

At each timestep, we compute the change in total $Ca^{2+}$ ($\Delta C_i$) and total binding sites ($\Delta S_i$) on each mesh cell, using standard finite-volume methods [16] to find the diffusive flux (via a vertex-centered difference scheme, detailed below) and advective flux (via a Lax-Wendroff scheme [16]) into or out of each mesh cell, in accordance with Eq. 2equation.0.2. Terminal boundaries of a mesh cell are taken to be no-flux boundaries. The change in total $Ca^{2+}$ due to leakage from each permeable cell includes an additional term $\Delta C_{\text{i,leak}} = -PA_iU_i\Delta t/V_i$. The current out of the network is given by $J(t) = -\sum_i V_i \frac{\Delta C_{\text{i,leak}}}{\Delta t}$, with the summation over all permeable mesh cells.

From the equilibration constraint (Eq. S2), we find the change in free $Ca^{2+}$ at each time-step, according to:

$$\Delta U_i = \frac{\Delta C_i - U_i \Delta S_i/(U_i + K_D)}{1 + S_i K_D/(U_i + K_D)^2} \tag{S6}$$

The concentration fields are then propagated forward in small time-steps $\Delta t = 2 \times 10^{-5}$s for purely diffusive transport and $\Delta t = 10^{-6}$s in the presence of flows. The current out of the network at each time-step is integrated over time to find total $Ca^{2+}$ released, or plugged into Eq. S5 to compute the cytoplasmic $Ca^{2+}$ level.

For active network simulations, flow velocities of magnitude $|v| = 20\mu$m/s are defined in a randomly chosen direction on each edge along the network. After each time step, the velocity along each edge flips direction with probability $1 - \exp(-\Delta t/\tau)$. Because each non-terminal mesh-cell boundary is placed along a network edge, the velocity across each boundary is uniquely defined and is used in the Lax-Wendroff calculation of the advective flux.

*Diffusive flux with enlarged well-mixed reservoirs*

For two mesh cells representing neighboring segments of tubule, the diffusive flux along the boundary between them can be written in the standard form:

$$\mathcal{F}^{(U)}_{i+1/2} = -D_{\text{Ca}} \frac{U_{i+1} - U_i}{\ell_{i+}} \tag{S7}$$

where $\ell_{i+} = (\Delta l_i + \Delta l_{i+1})/2$ is the separation between mesh cell centers. An analogous expression can be written for $\mathcal{F}^{(B)}_{i+1/2}$ corresponding to the flux of bound $Ca^{2+}$ (defined as $B_i = C_i - U_i$), with $D_{\text{Ca}}$ replaced by $D_b$. The change in $Ca^{2+}$ concentration due to diffusive transport is then given by:

$$\frac{\Delta C_{i,\text{diff}}}{\Delta t} = \frac{\pi r^2}{V_i}\left(\mathcal{F}^{(U)}_{i+1/2} - \mathcal{F}^{(U)}_{i-1/2} + \mathcal{F}^{(B)}_{i+1/2} - \mathcal{F}^{(B)}_{i-1/2}\right). \tag{S8}$$

For the case where one of the mesh cells represents a large well-mixed reservoir, the expression for the diffusive flux must be altered to account for the narrow escape process of diffusive particles leaving the reservoir through small tubule entrances. For the partially fragmented ER network structure in RTN3 OE cells, the individual bubble-like fragments are treated as spheres with surface area $A_{\text{bub}} = 4\pi R^2_{\text{bub}}$ and volume $V_{\text{bub}} = \frac{4}{3}\pi R^3_{\text{bub}}$.

For such a spherical reservoir connected to a narrow tubule of radius $r$, we consider the steady state flux from the sphere into the tubule, where we assume the concentration throughout the sphere is $U_{\text{bub}}$ and the concentration in the tubular mesh cell at a distance $\Delta l_i/2$ from the sphere is $U_i$. Defining $\tilde{U}_i$ as the concentration at the sphere-tubule junction point, the current out of the sphere must be $I_{\text{bub}} = 4Dr(U_{\text{bub}} - \tilde{U}_i)$ [17]. The current from the tube into the sphere is $I_{\text{tube}} = D\pi r^2(\tilde{U}_i - U_i)/(\Delta l_i/2)$. Setting these two expressions equal to each other at steady-state gives the flux across the boundary between the two mesh cells as:

$$\mathcal{F}^{(U)}_{\text{bub},i} = \frac{4D(U_{\text{bub}} - U_i)}{\pi r + 2\Delta l_i}. \tag{S9}$$

This flux expression can be put in the same form as the boundary between two tubular cells (Eq. S7), if we take the reservoir to have an effective length $\Delta l_{\text{bub}} = \pi r/2$. This effective length does not depend on the size of the sphere, because diffusive transport to a narrow target in 3D is non-compact [18] and the current of escaping particles is thus independent of the size of the confining domain.

To propagate forward the concentration fields in the spherical reservoir, we use an expression analogous to Eq. S8, using the reservoir volume $V_{\text{bub}}$, and incorporating flux across all boundaries to connected tubules.

#### Cell culture, transfections and constructs and reagents

The constructs used are listed in the supplemental plasmid list (Table S1) and will be available on Addgene. COS-7 (RRID:CVCL_0224) and EGFP-IP3R1 HeLa ([19]) cells were cultured in Dulbecco's modified Eagle's medium (DMEM) only or DMEM/F12 respectively, supplemented with 10% fetal bovine serum (FBS), 2 mM L-Glutamine and 100 U/mL Penicillin-Streptomycin (P/S). Cells were transfected using the Neon Transfection System (Invitrogen) with 2 pulses of 20ms at 900V for COS-7 cells and 2 pulses of 35ms at 1,005V for HeLa cells. RTN3 OE cells were obtained by transfecting COS-7 cells with RTN3-HaloTag plasmid monitored by labelling with JF646; ATL2/3 KO cells were obtained from Junjie Hu (University of Chinese Academy of Sciences).

Caged-$IP_3$ (ci-$IP_3$/PM, #6210) was obtained from Tocris. Thapsigargin (10522-1 mg-CAY) was obtained from Cambridge Bioscience (Cambridge, UK). Janelia Fluor 646-conjugated HaloTag ligand (JF646, not photoactivatable, GA1120) were obtained from Promega (Madison, WI, USA).

#### Continuous photoactivation chase (CPAC)

##### *Image acquisition*

This assay was performed as previously described in [8]. Briefly, COS-7 cells were transfected with $\text{paGFP}^{\text{ER}}$ plasmid and $\text{mCherry}^{\text{ER}}$ plasmids (together with RTN3-HaloTag plasmid for RTN3 OE cell line) and 48h post-transfection, cells were taken to microscopy. We used a confocal microscope (STELLARIS8, Leica, Wetzlar, Germany) with a controlled environment (37°C, 5% $CO_2$) and the following parameters: excitation/emission of 489 nm/$495 - 550$ nm for $\text{paGFP}^{\text{ER}}$ and 567 nm/$608 - 635$ nm emission for $\text{mCherry}^{\text{ER}}$, frame size of $512 \times 512$ pixels, scan speed of 1000 Hz, 1.92 frames per second.

Image acquisition series consisted of pre-photoactivation images (10 frames), photoactivation illumination (405 nm, 100% laser power) introduced in a region of interest (photoactivation spot of $5\mu\text{m} \times 5\mu\text{m}$) for a duration of at least 300 frames (520 ms/frame), using the Fly mode (enabling simultaneous photoactivation and image recording) in the FRAP wizard. Image series were then analyzed semi-automatically as described in the following section.

##### *Image analysis*

$\text{paGFP}^{\text{ER}}$ mobility in the ER lumen was determined by photoactivation image analysis and median rise-time estimation using a custom code written in Matlab (version 9.5.0, R2018b, Natick, Massachusetts: The MathWorks Inc.), as described in Ref. [8]. Briefly, the images of ER luminal marker $\text{mCherry}^{\text{ER}}$ were used to segment the cell and nucleus shapes manually. The resulting ROI consisted of the area within cell boundary except for the nucleus (eroded by 1 micron to avoid edge effects). The photoactivation region was manually segmented from images of $paGFP^{ER}$ focused on the activation spot during the photoactivation phase and removed from the cell ROI to be analyzed (with a 1$\mu$m buffer). Concentric ring regions of width 2 $\mu$m were defined around the photoactivation center (centroid of the photoactivation region), starting at 1 $\mu$m intervals in distance from it, and sub-divided into wedge-shaped ROIs of arc length 2 $\mu$m, shifted by 1 $\mu$m around the circle. Wedge ROIs presenting less than 2 $\mu m^2$ of valid area after intersection with the segmented cell area were not retained for analysis.

Fluorescence intensity time traces were obtained from every wedge-shaped ROI and normalized by the ROI area. The intensity time series were all fit to a double-exponential function beginning after the photoactivation start and rising from the pre-photoactivation value to a maximum value set by the final fluorescence of the photoactivated region (assuming that fluorescence in the whole cell will equilibrate to this value at infinite time). The exponential fits served to compute half-time for the signal rise, unless signal-to-noise ratio was insufficient (signal range less than twice the pre-photoactivation standard deviation) or resulting half-time was more than 10 times the recorded timespan. Median arrival $t_{1/2}$ corresponds to the median half-time of all remaining wedge regions at a given radius from the photoactivation center, over all cells in each dataset. The error bars (standard deviation of the median) were obtained by bootstrapping individual cells from each dataset and repeating the analysis to indicate cell-to-cell variation in each reported half-time.

The code for analysing CPAC measurements is available on `https://github.com/lenafabr/photoActivationAnalysis.git`

**Fluorescence Lifetime Imaging Microscopy (FLIM)**

To measure $[Ca^{2+}]_{ER}$, we used Fluorescence Lifetime Imaging Microscopy (FLIM) and a probe inspired from D4ER FRET-FLIM based probe (ECFP/Citrine pair [20]). We improved this probe by swapping the donor fluorophore ECFP to mTurquoise and acceptor Citrine to REAch2. These modifications were expected to provide more accurate $[Ca^{2+}]_{ER}$ estimations as mTurquoise decay can reliably be fitted with a monoexponential as opposed to ECFP and REAch2 is a dark acceptor (nonfluorescent yellow fluorescent protein (YFP) mutant). We performed a calibration of the new version of D4ER, named D4ER-Tq, by inducing maximum $Ca^{2+}$ condition with addition of ionomycin (10$\mu$M) and minimal $Ca^{2+}$ condition was obtained by adding thapsigargin (3$\mu$M) to trigger full depletion of the $Ca^{2+}$ stored in the ER. The dynamic range of this new version of D4ER was increased from 0.8ns to 1ns. This maximal and minimal values were used to convert lifetimes to $Ca^{2+}$ concentrations. COS-7 cells (WT, RTN3 OE monitored by labelling with JF646, ATL2/3 KO, mCherry-tagged Climp63 OE lines) were transfected to express D4ER-Tq probe and 48h post-transfection, cells were taken to microscopy. We used a confocal microscope (STELLARIS8, Leica, Wetzlar, Germany) with a controlled environment (37°C, 5% $CO_2$) and the following FLIM parameters: excitation/emission of $440/450-500$ nm, pulsed wight light laser at 80 MHz, HyDX detector in counting mode, frame size of $512\times512$ pixels, scan speed of 400 Hz, and settings were set to reach 5000 photons/pixel. Images were processed using Leica STELLARIS8 FLIM wizard. ROIs were drawn around individual cells and data fitted to a monoexponential decay function. Lifetime values were converted to $[Ca^{2+}]_{ER}$ using the following formula, with D4ER Kd (321 $\mu$M), Hill coefficient (1.01) and the $\tau_{max}$ and $\tau_{min}$ determined during calibration as reported in [20]:

$$[Ca^{2+}] \approx K_D\left(\frac{\tau_{\rm max}-\tau_{\rm exp}}{\tau_{\rm exp}-\tau_{\rm min}}\right)^{1/h}. \tag{S10}$$

##### Measurement of total $Ca^{2+}$ load

To measure total $Ca^{2+}$ load in the ER lumen, COS-7 cells (WT, ATL2/3 KO, mCherry-tagged Climp63 OE) were transfected to express GCaMP3 probe, an ER membrane-tethered $Ca^{2+}$ sensor facing cytosolic side [21] or incubated with Oregon Green BAPTA-1, AM at 5$\mu$M for 30 min (COS-7 RTN3 OE monitored by labelling with JF646). 48h post-transfection, cells were taken to microscopy. Cells were treated with thapsigargin (1 $\mu$M) to inhibit the SERCA pump and deplete ER $Ca^{2+}$. We used a confocal microscope (STELLARIS8, Leica, Wetzlar, Germany) with a controlled environment (37°C, 5% $CO_2$) with excitation/emission of $480/485-550$ nm with 0.1 frames per second for 10 minutes post treatment. Peaks indices and lowest contour lines were automatically detected and used to quantify the Area Under the Peak with the composite trapezoidal rule in Python.

##### Local $Ca^{2+}$ release events generation, imaging, and analysis

*Image acquisition*

COS-7 cells (WT, RTN3 OE monitored by labelling with JF646, ATL2/3 KO, mCherry-tagged Climp63 OE lines) were transfected to express GCaMP3 probe, and 48h post-transfection, cells were taken to microscopy. Local release was achieved by treating cells with caged-$IP_3$ (3 $\mu$M, 2.5 hours) followed by low amounts of Bapta-AM (30nM, 5min), a cell permeant chelator. $Ca^{2+}$ chelation in the cytoplasm is employed to limit the spread of $Ca^{2+}$ ions and to prevent local releases from expanding into global events via $Ca^{2+}$-induced $Ca^{2+}$ release [4, 22, 23]. We used a confocal microscope (STELLARIS8, Leica, Wetzlar, Germany) with a controlled environment (37°C, 5% $CO_2$) and the following parameters: excitation/emission of $480/485-550$ nm, frame size of $512\times256$ pixels, scan speed of 800 Hz bidirectional, 3 frames per second. Image acquisition series consisted of pre-photo-uncaging images (50 frames), photo-uncaging of caged-$IP_3$ achieved by short illumination (405 nm, 50% laser power) across the whole cell using the Fly mode in the FRAP wizard (time interval 150-330ms), and post-photo-uncaging images (150 frames). We confirmed that $IP_3$R were present and functional in all cell types as we could trigger global $Ca^{2+}$ release through CICR [24] upon uncaging high amounts of $IP_3$ with long laser exposure times (30s). The 405nm laser pulse duration was then reduced to trigger local releases only, while avoiding the induction of global $Ca^{2+}$ waves across the cytoplasm.

Local releases were then analyzed semi-automatically as described in the following section.

*Image analysis*

The magnitude of local $Ca^{2+}$ transients was determined via image analysis using a custom code written in Matlab [25]. The images of $Ca^{2+}$ sensor GCaMP3 during post-photo-uncaging image series were used. Cell shape was manually segmented and the resulting ROI consisted of the area within cell boundary. This ROI was sub-divided by tiling the cell area into a set of rectangular regions of size $25 \times 25$ pixels. Signal traces from each rectangular sub-region were plotted and used to search for peaks in the resulting signal traces. Filtering for peaks was achieved by detrending the signal with polynomial fit (order 3) followed by SVG smoothing (order 5, window size 15).

Signal peaks were identified by thresholding using a universal threshold based on the median absolute deviation of the detrended signal [26]. In addition, the detrended signal curvature was computed at each peak, and only those peaks with negative curvature beyond a threshold value (set to $0.1\times$ the universal threshold) were retained for further analysis. ROIs with similar peak positions in time (according to a set cutoff for time separation) and adjacent in space were combined together to prevent similar $Ca^{2+}$ signals to be measured through several ROIs. The peak $Ca^{2+}$ signal within the merged cluster of regions was identified and used to create a new rectangular ROI centered on that peak ($35 \times 35$ pixels). Signal traces were extracted from these new ROIs and peak detection again applied, using the same threshold values. A total of 59, 29 and 47 peaks were analyzed for WT, RTN3 OE and ATL KO respectively. The integral of the area under the peak was found by approximating each peak as a Gaussian shape of a fitted width as given by Matlab's 'findpeaks' function. The peak integrals were used as a measure of the magnitude of $Ca^{2+}$ release in the ROI.

The code for analysing local $Ca^{2+}$ releases is available on `https://github.com/lenafabr/photoActivationAnalysis/roisignalGUI.git`

##### 3D Fluorescence microscopy of $IP_3Rs$

Prior to 3D fluorescence imaging, EGFP-IP3R1 HeLa cells were transfected to express $mCherry^{ER}$ and RTN3 monitored by labelling with JF646. We used a confocal microscope (STELLARIS8, Leica, Wetzlar, Germany) with a controlled environment (37°C, 5% $CO_2$) and the following parameters: excitation/emission of $488/500-550$ nm for EGFP-IP3R1, of $555/575-640$ nm for $mCherry^{ER}$, of $646/670-750$ nm for JF646, frame size of $1024 \times 1024$ pixels, scan speed of 800 Hz unidirectional, z-step of $0.29\mu$m. Images were post-processed using Leica Lightning deconvolution and analyzed using 3D Objects Counter and Distance Analysis (DiAna) plugins in Fiji to determine cluster centre of mass from 3D z-stacks and measure distance to closest neighbour respectively.

##### Single-step photobleaching of $IP_3Rs$

Prior to imaging, EGFP-IP3R1 HeLa cells were transfected to express $Halo^{ER}$ and RTN3-mScarlet3 and fixed with 2% PFA, 2% glutaraldehyde, 100 mM cacodylate (pH 7.4), and 2 mM CaCl2 for 30 min at room temperature (RT). TIRF time-lapse images were acquired (EGFP, 488nm; mScarlet3, 561nm; HaloER, 647nm) at 10 fps for 90 seconds.

The TrackMate plugin in Fiji was used to detect EGFP-IP3R1 puncta and extract puncta density (together with cell area measurement) and intensity traces over time. The quickPBSA package in Python was used to determine the number of photobleaching steps in each trace ([27]).

**Supplemental Plasmid List**

| ID | Plasmid name | Description | Reference | First appearance | Label in figure |
|---|---|---|---|---|---|
| 56 | pLV_TRE3G_hRTN3E-Halo | Tet-inducible hRTN3E (long form) expression vector, expresses hRTN3E-TEV-Halo, Lentiviral vector, from VectorBuilder | this paper | Figure 3 | RTN3 OE |
| 55 | pmPA-GFP-ER-5 | Mammalian expression of ER-targeted photoactivatable GFP | Addgene #57132 | Figure 3 | paGFPER |
| 230 | pLV_Tre3g_ CalcireticulinSS_ mTurquoise_ D4-cameleon_ REAch2_KDEL | FRET-based calcium sensor located in the ER lumen | this paper | Figure 5 | D4ER-Tq |
| 16 | pcDNA3-ER-GCaMP3 | Mammalian expression, ER-targeted $Ca^{2+}$ indicator GCaMP3; targeted to cytosolic face of the ER membrane | PMID:25056880 Addgene #64854 | Figure 5 | GCaMP3 |
| 146 | pAAV_RTN3E-Halo | CAG promoter-regulated human RTN3E expression vector (adenoviral backbone) | this paper | Figure S12 | RTN3-Halo |
| 701 | pCMV_RTN3E-mScarlet3 | Mammalian expression vector for RTN3E-mScarlet3 | this paper | Figure S13 | RTN3-mScarlet3 |
| 681 | pLV_CMV_Prss-Halo-KDEL | CMV promoter-regulated ER targeted HaloTag expression vector | this paper | Figure S13 | HaloER |
|  | mCh-Climp63 | CMV promoter-regulated expression of Climp63, pmCherry-N1 backbone | Addgene #136293 | Figure S11 | Climp63 |

TABLE S2. List of plasmids used in this study

#### S2. SUPPLEMENTAL RESULTS

##### Cytoplasmic buffer proteins in local release model

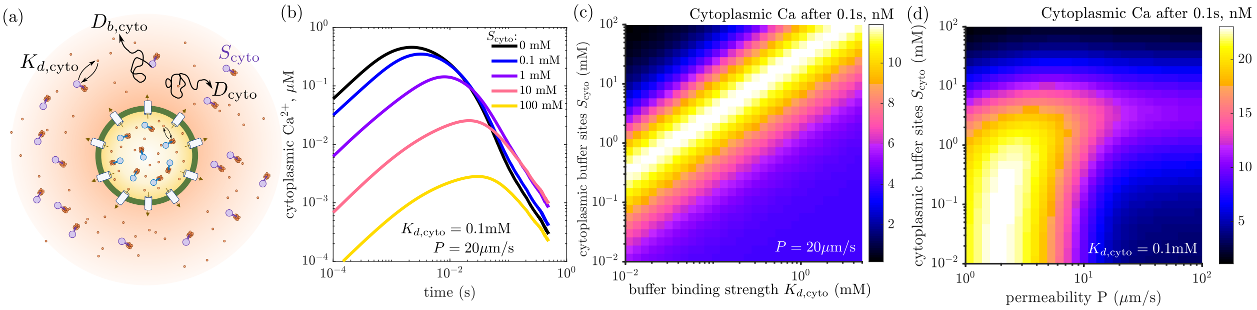


FIG. S1. Cytoplasmic $Ca^{2+}$ levels with inclusion of cytoplasmic buffer binding, for model with release of local $Ca^{2+}$ pool only (no luminal transport). (a) Model schematic, indicating equilibrated binding to diffusive buffer sites in the cytoplasm. (b) Cytoplasmic $Ca^{2+}$ concentration (in $\mu$M) near release site over time, for several different densities of cytoplasmic binding sites $S_{\rm cyto}$. (c) Averaged free cytoplasmic $Ca^{2+}$ level (in nM) near release site, as a function of cytoplasmic buffer site density $S_{\rm cyto}$ and cytoplasmic buffer strength $K_{d,\rm cyto}$. (d) Averaged free cytoplasmic $Ca^{2+}$ near release site as a function of buffer site density and permeability of local release region $P$. $Ca^{2+}$ concentrations are shown at time $t = 0.1$ sec, as an average over a sphere of radius $R_{\rm loc} = 1\mu$m, centered on the release site. Total length of releasing tubules is set to $L = 0.75\mu$m, cytoplasmic $Ca^{2+}$ diffusivity is $D_{\rm cyto} = 200\mu{\rm m}^2$/s, and cytoplasmic buffer diffusivity is $D_{b,\rm cyto} = 2\mu{\rm m}^2$/s.

The levels of free $Ca^{2+}$ in the cytoplasm surrounding the release site can be modulated by binding of $Ca^{2+}$ to buffer proteins. Because estimates for the density and binding strength of cytoplasmic $Ca^{2+}$ buffers vary widely, we consider a range of values here, with the goal of demonstrating that the presence of buffers cannot substantially increase the local concentration of free $Ca^{2+}$.

We incorporate cytoplasmic buffers into our model for purely local release numerically, using a finite-volume approach in radial coordinates and assuming equilibrated binding. The unbound $Ca^{2+}$ density $\psi_u(\rho, t)$ and bound $Ca^{2+}$ density $\psi_b(\rho, t)$ (as a function of radial distance $\rho$ from the release site) obey the following dynamic equation:

$$\frac{d\psi}{dt} = D_{\rm cyto}\nabla^2\psi_u + D_{\rm b,cyto}\nabla^2\psi_b + J(t)\frac{\delta(\rho)}{4\pi\rho^2}, \tag{S11}$$

where $D_{\rm cyto} = 200\mu{\rm m}^2/{\rm s}$ and $D_{b,\rm cyto} = 2\mu{\rm m}^2/{\rm s}$ are diffusivities of free $Ca^{2+}$ and buffer proteins, respectively, $J(t)$ is the release current, and $\delta$ is the Dirac delta function. The equilibrium condition is expressed as

$$K_{D,\rm cyto} = \frac{\psi_u(S_{\rm cyto} - \psi_b)}{\psi_b}, \tag{S12}$$

where $K_{D,\rm cyto}$ is the cytoplasmic buffer binding strength and $S_{\rm cyto}$ is the (constant) density of cytoplasmic buffer binding sites.

To evolve forward Eq. S11, we break up the space surrounding the origin into mesh cells representing spherical shells, of thickness $\Delta\rho$, and volume $V_i = 4\pi\rho_i^2\Delta\rho$, for $i = 1, 2, \ldots$. The innermost mesh cell has volume $V_0 = \frac{4}{3}\pi(\Delta\rho)^3$. The current of free $Ca^{2+}$ through the inner and outer boundary of mesh cell $i$ is then

$$\mathcal{I}^u_{i-1/2} = D_{\text{cyto}} 4\pi \left(\rho_i - \frac{\Delta\rho}{2}\right)^2 \frac{(\psi_{u,i} - \psi_{u,i-1})}{\Delta\rho}$$
$$\mathcal{I}^u_{i+1/2} = D_{\text{cyto}} 4\pi \left(\rho_i + \frac{\Delta\rho}{2}\right)^2 \frac{(\psi_{u,i+1} - \psi_{u,i})}{\Delta\rho}, \tag{S13}$$

with analogous expressions for the diffusive current of bound $Ca^{2+}$. At each time-step, we compute the change in total $Ca^{2+}$ in each mesh cell due to diffusive current, analogous to Eq. S8 but with the area of each boundary already incorporated into the current. Additionally, the innermost cell ($i = 0$) concentration is updated due to the $Ca^{2+}$ release according to $J(t)\Delta t/V_0$. The total $Ca^{2+}$ field is evolved forward in time-steps of $\Delta t = 10^{-5}$s, and the unbound and bound $Ca^{2+}$ fields are concurrently updated via Eq. S12.

The cytoplasmic free $Ca^{2+}$ concentration is averaged over a sphere of radius $R_{\text{loc}} = 1\mu$m and plotted versus time in Fig. S1b. Increasing the buffer concentration slows the spread of $Ca^{2+}$ from the release site, but also decreases the fraction of cytoplasmic $Ca^{2+}$ that is free from buffer binding. Parameter sweeps for different values of membrane permeability $P$ and cytoplasmic buffer binding strength $K_{d,\text{cyto}}$ indicate that the free $Ca^{2+}$ concentration around the release site at 0.1s never rises above 30nM, regardless of how many cytoplasmic buffer sites are present (Fig. S1c,d). Thus, cytoplasmic buffering does not negate the need for delivering $Ca^{2+}$ through the lumen to supplement the depleted local release region in order to enable high cytoplasmic $Ca^{2+}$ levels.

#### Luminal buffer carrying capacity

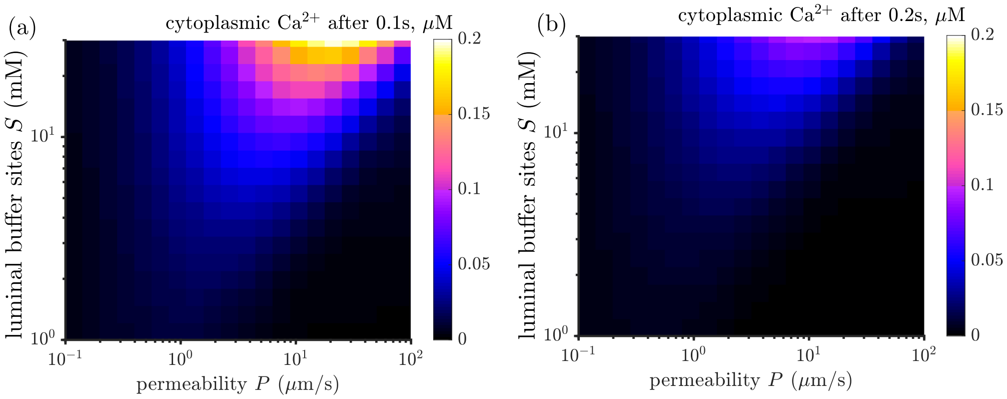


FIG. S2. Cytoplasmic $Ca^{2+}$ levels from purely local release vary with luminal $Ca^{2+}$ carrying capacity. (a) Cytoplasmic $Ca^{2+}$ concentration averaged over a region of radius $R_{\text{loc}} = 1\mu$m, at time $t = 0.1$sec, plotted as a function of membrane permeability $P$ and concentration of luminal binding sites $S$. (b) Corresponding plot for local cytoplasmic $Ca^{2+}$ at $t = 0.2$sec.

Throughout the calculations in the main text, we make a conservative estimate of the luminal $Ca^{2+}$ carrying capacity. Namely, we assume that buffer protein concentration is $\sim 0.1$mM and that each protein contains 25 $Ca^{2+}$ binding sites ($S = 2.7$mM). Given the presence of multiple luminal $Ca^{2+}$ buffers, the actual capacity is likely to be higher. As a consequence, purely local release from an isolated region of size $R = 0.25\mu$m will result in higher cytoplasmic $Ca^{2+}$ concentrations with higher values of $S$ (Fig. S2). However, we note that even an order-of-magnitude increase in

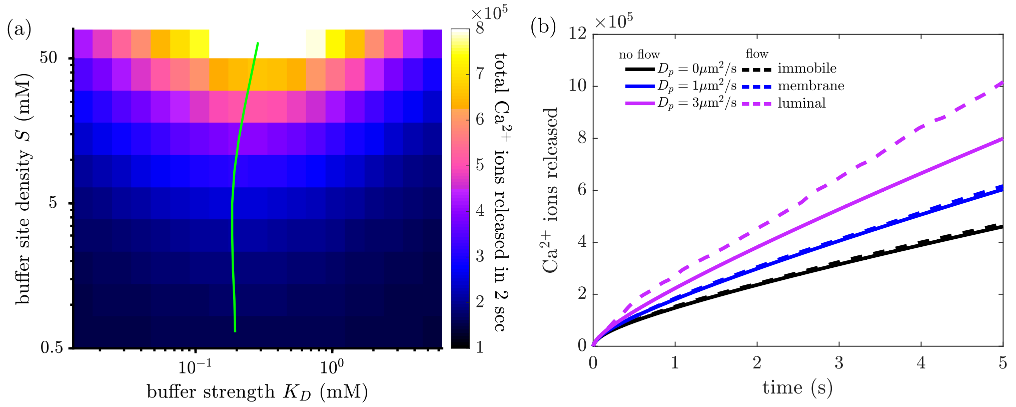


FIG. S3. $Ca^{2+}$ release from honeycomb network with increased luminal buffer site concentration. (a) Cumulative $Ca^{2+}$ release in 2sec, as a function of buffer binding strength $K_D$ and concentration of luminal buffer sites $S$. Green line shows optimal binding strength for each value of $S$. (b) Cumulative $Ca^{2+}$ release over time for buffer proteins with different diffusivities, with (dashed lines) and without (solid lines) ER luminal flows. Results shown assume a buffer site density of $S = 10$mM. Compare to Fig. 2h, which uses $S = 2.5$mM.

buffer sites (to $S = 25$mM), will result in cytoplasmic $Ca^{2+}$ concentrations below 200nM at 0.1sec after the start of release. These concentrations drop still further at subsequent times. We therefore maintain that purely local release is insufficient to maintain the observed magnitudes of $Ca^{2+}$ puffs (which can range up to $1\mu$M in osteoclasts [28]), even in the case where the luminal $Ca^{2+}$ capacity is substantially higher.

For the spatial model incorporating transport through the ER network, the results presented in this work become even more pronounced if there is a higher concentration of luminal buffer sites. Specifically, the enhancement in $Ca^{2+}$ release at the optimal binding strength $K_d$ is greater with increasing buffer capacity (Fig. S3a). The value of that optimal strength remains at $K_d^{\text{opt}} \approx 0.2$mM over a broad range of buffer concentrations. Furthermore, the effect of luminal buffer protein mobility, both with and without active flow within the ER tubules, is augmented when there are more buffer proteins present (Fig. S3b, compare to Fig. 2h).

#### Estimate of luminal protein diffusivity

In our $Ca^{2+}$ dynamics simulations, we take $D_b = 2.8\mu m^2/s$ as an estimate of ER luminal protein diffusivity. This estimate was obtained from single-particle trajectory analysis and falls within the range of published estimates, which indicate that diffusion in the lumen may be reduced by 2-10 fold compared to the cytoplasm [29–31].

Here we validate this estimate by analysis of the spatiotemporal dynamics of $\text{paGFP}^{\text{ER}}$ (as in Fig. 3), compared to simulations of diffusive particle spread on extracted ER network structures.

Specifically, we use the same finite volume method employed throughout the manuscript to simulate network diffusion of a single species (no binding) with $D = 2.8\mu m^2/s$. A circular region of radius $1.4\mu$m is set to a fixed value of 1 (arbitrary units), in different locations of the network, representing the photoactivated region. The concentration profile around this photoactivated region is then computed over time (Fig. S4a). This profile is blurred with a Gaussian filter of radius $\sigma = 1\mu$m and the resulting images for each timepoint are processed in the same way as the experimental data shown in Fig. 3. Signal is tracked in many small ROIs at different distances from the photoactivated

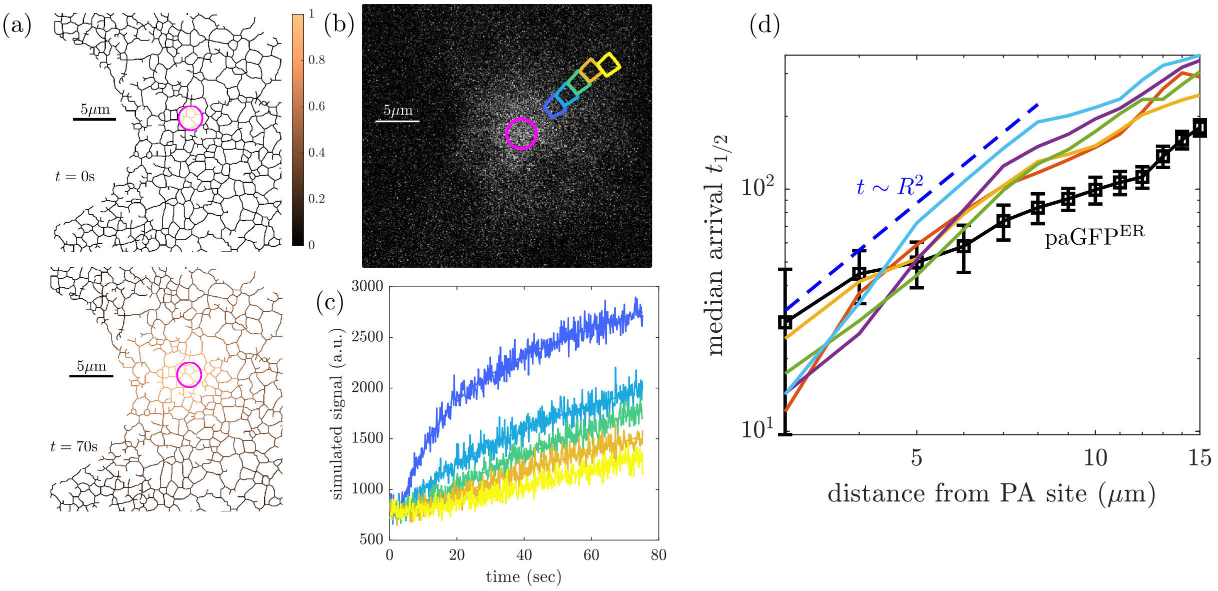


FIG. S4. Simulation of photoactivated protein spreading through an ER network. (a) Snapshots of concentration profiles for the spreading of particles with diffusivity $D = 2.8\mu\text{m}^2/\text{s}$, at time $t = 0$sec and $t = 70$sec. The photoactivated region (magenta) is held at a fixed unit concentration. (b) Blurred concentration profile analyzed analogously to experimental data, with signal in different wedge-shaped regions tracked over time. (c) Temporal evolution of signal in the colored ROIs marked in part (b). (d) Median arrival half-time for all wedge ROIs at a given distance from the photoactivation center, plotted on log-log axes to show scaling. Black: experimental data, as in Fig. 3c. Each solid colored line corresponds to simulations with 5 different photoactivated regions, on a network structure extracted from a single cell. Dashed blue line shows quadratic scaling expected for diffusive particles.

center (Fig. S4b,c) and fit to a double-exponential function to extract an effective half-time $t_{1/2}$ for signal arrival.

Plotting the median arrival half-time versus the distance from the photoactivated center for simulated data shows the expected diffusive scaling ($t_{1/2} \sim R^2$, Fig. S4d). By contrast, the experimental data for paGFP$^{ER}$ exhibits a super-diffusive scaling as previously reported [8]. This scaling is consistent with the notion that there may be some active mechanisms (such as luminal flows) that enhance the ability of luminal contents to spread over larger distances. The mismatch between experimental photoactivated spreading data and theoretical predictions of diffusive scaling justifies a variation of the model with random active flows driving luminal contents, as shown in Fig. 2c-h. Such active flows further increase the importance of mobile (rather than membrane-bound) buffer proteins in enhancing the magnitude of $Ca^{2+}$ release.

Throughout the rest of the manuscript, we rely on the simpler, more-established, conventional model for intra-ER transport and assume purely diffusive motion of buffer proteins and free ions. On length scales below approximately $7\mu$m, the simulated diffusive arrival times for particles with $D = 2.8\mu\text{m}^2/\text{s}$ approximately match the experimental observations. Therefore, this value of the diffusivity serves as a good approximation for the actual spreading rates of proteins within the ER lumen. It is taken as the luminal protein diffusion coefficient throughout our manuscript.

The diffusivity of $Ca^{2+}$ ions in the lumen cannot be readily measured, so we estimate it to be a factor of 10 higher than the protein diffusivity [9] and set $D_{\text{Ca}} \approx 28\mu\text{m}^2/\text{s}$.

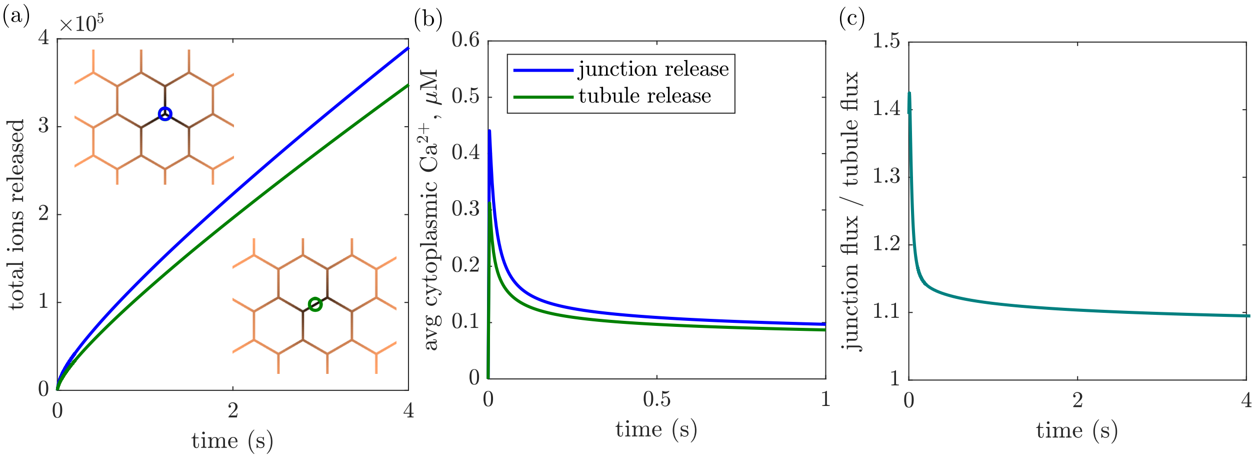


FIG. S5. Release events from tubule versus junction regions have similar magnitude. (a) Cumulative $Ca^{2+}$ ion released over time in a honeycomb network with the release region (radius $R = 0.25$) centered on a junction (blue) or the middle of a tubule (green). Insets show snapshot near release region at time 2s. (b) Cytoplasmic $Ca^{2+}$ concentration averaged over a region of radius $R_{\text{loc}} = 1\mu$m, plotted over time for same network release centers as in (a). (c) Ratio of $Ca^{2+}$ flux out of the network over time, between release centered at a junction and release centered in the middle of a tubule.

#### Release from tubules vs junctions

The positioning of the release region within a network structure can potentially affect the magnitude of the release. In particular, one might expect that centering the release region on a degree-3 junction node would lead to greater release than the same-size region centered on a degree-2 position along the tubule. As shown in Fig. S5, this is indeed the case for very short times ($< 0.1$s). When release occurs at a junction, there is 50% more length of permeable tubule capable of releasing ions, leading to a 50% higher flux at short times. However, for times beyond this initial transient period, the difference between flux from the two different release zones is only about 10% (Fig. S5c). This observation highlights the importance of transport to the release region in limiting the efflux of ions. After the release region itself is depleted, this transport becomes rate-limiting and the flux is no longer simply proportional to the length of permeable tubule.

#### Optimal buffer strength for different permeabilities

In the model incorporating $Ca^{2+}$ and buffer transport through a network, the optimum buffer binding strength is robust over a broad range of time-scales (Fig. 1e). However, this optimal binding strength does depend on the permeability $P$ and extent $R$ of the local release region. When the leaking of $Ca^{2+}$ from the release region is much slower than its diffusive delivery from elsewhere in the lumen, the local free $Ca^{2+}$ concentration is maintained close to its original value of $U_0 = 0.5$mM, and the optimal binding strength $K_D^{\text{opt}} \rightarrow 0.5$mM concomitantly (Fig. S6). By contrast, when local release is much faster than diffusive delivery, the local $Ca^{2+}$ concentration drops and $K_D^{\text{opt}}$ becomes smaller, in order to maximize the total amount of $Ca^{2+}$ ions stored in the lumen.

The fully connected honeycomb network considered here can be approximately treated as an effective two-dimensional continuum, with the initial free $Ca^{2+}$ concentration per unit area given by $\widehat{U}_{2D} = U\pi r^2\rho$, where $U$ is the concentration of free $Ca^{2+}$ in the lumen and $\rho$ is the density of ER tubule length per unit area. For a honeycomb network with edges of length $\ell$, this density is

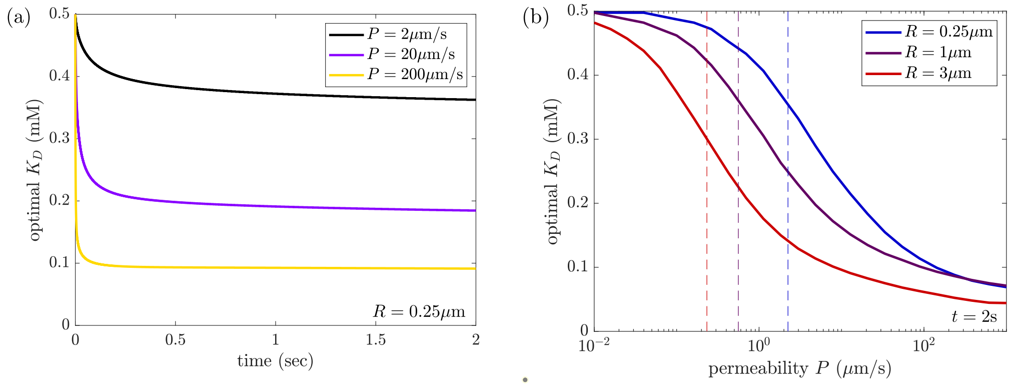


FIG. S6. Effect of permeability and release region size on optimal buffer strength. (a) Value of binding strength $K_D$ that maximizes the cumulative amount of $Ca^{2+}$ released by a certain time, plotted as a function of time. Each curve corresponds to a different permeability of the release region. The optimal $K_D$ is robust over a broad range of times. (b) Optimal binding strength $K_D$ that maximizes cumulative $Ca^{2+}$ release in 2s, plotted versus the permeability $P$. Each curve corresponds to a different radius of the release region ($R$). Dashed lines show estimated critical permeability $P^*$ beyond which $Ca^{2+}$ transport becomes rate-limiting and local $Ca^{2+}$ levels are depleted. Below $P^*$, the optimal binding strength matches the initial free $Ca^{2+}$ ($U_0 = 0.5$mM). Above $P^*$, the optimal $K_D$ drops to lower concentrations.

$\rho = 2/(\sqrt{3}\ell)$. To estimate the rate of arrival to the release region, we use the long-time limit for two-dimensional diffusive current to a region of size $R_{\text{eff}}$, assuming an effective diffusivity $D/2$ [32]:

$$k_{\text{2D}} \approx \frac{2\pi D \widehat{U}_{2D}}{\log(2Dt/R_{\text{eff}}^2) - 2\gamma}, \tag{S14}$$

where $\gamma \approx 0.58$ is the Euler–Mascheroni constant. This expression takes into account the factor of 2 reduction in effective 2D diffusivity due to confinement in a lattice of tubules [33]. The effective capture radius is set to $R_{\text{eff}} = \max(R, \ell)$, corresponding to the distance where the tubules leading to a small release region join the rest of the lattice, or else to the size of an entire larger release region. The logarithmic term implies that the diffusive delivery (and hence the local $Ca^{2+}$ concentration) is not very sensitive to time, and helps explain the lack of time-sensitivity for $K_D^{\text{opt}}$.

The local release current from a permeable region with total tube length $L$ is given by $k_{\text{release}} = P(2\pi r)L$. As shown in Fig. S6b, the optimum binding strength $K_D^{\text{opt}}$ transitions from $U_0$ down to 0mM when the permeability passes a critical value $P^*$. This value corresponds to the transition when release rates exceed diffusive delivery rates: $k_{\text{release}} > k_{\text{2D}}$, which corresponds to

$$P \gg P^* = \frac{\pi D r \rho}{L(\log(2Dt/R_{\text{eff}}^2) - 2\gamma)}. \tag{S15}$$

Here $L$ is the length of tubule in the releasing region ($L = 3R$ for $R < \ell$ and $L = \pi R^2 \rho$ for $R \gg \ell$). Notably, for large release regions, the critical permeability is independent of tubule density as the lattice behaves simply as an effective 2D continuum.

##### Optimal persistence time for transport in active network

An alternative model to purely diffusive transport for ER luminal content features an active network wherein particles undergo directed motion along individual edges [34]. While fluid flows

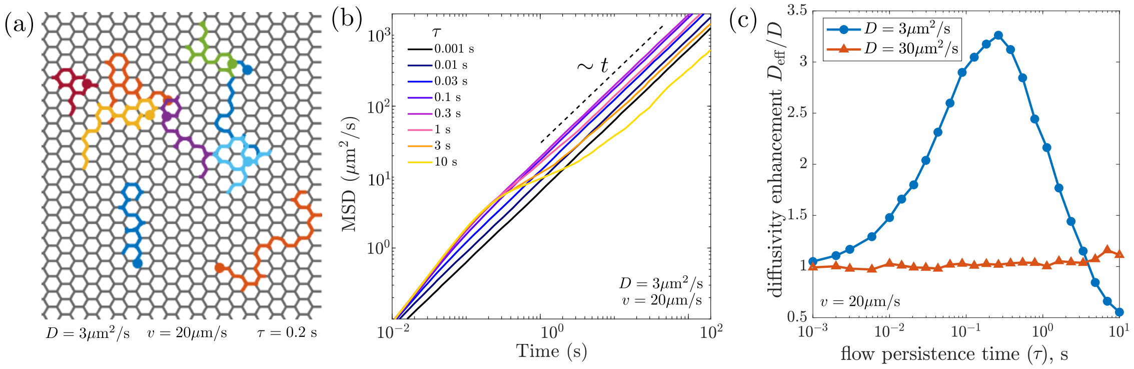


FIG. S7. Particle transport on network with active directed motion along edges. (a) Example trajectories of individual simulated particles on a honeycomb network with active flows along edges. Flow direction persists for average time $\tau = 0.2$s. Particles move by a combination of diffusion ($D = 3\mu\text{m}^2/\text{s}$) and drive $v = 20\mu\text{m/s}$. (b) Mean-squared displacement of simulated particles on active network, plotted for different values of the persistence time $\tau$. (c) Scaled effective long-time diffusivity $D_{\text{eff}}$, as a function of the active flow persistence time. Blue circles are simulations of particles with diffusivity $D = 3\mu\text{m}^2/\text{s}$; red triangles are particles with diffusivity $D = 30\mu\text{m}^2/\text{s}$. Edge flow velocity is $v = 20\mu\text{m/s}$ in both cases.

within the ER lumen have been proposed to account for the observed superdiffusive motion of luminal particles [13], the mechanism driving this motion remains unclear. Regardless of the underlying mechanism, we consider here the effect on transport of an active model with randomly flipping directed motion along the edges. Specifically, we assign to each edge a velocity $v = 20\mu\text{m/s}$, which flips direction independently on each edge as a Poisson process with timescale $\tau$.

For sufficiently long times ($t \gg \tau$), individual particles on such an active network undergo random walks that give rise to an effectively diffusive motion. We leverage agent-based simulations to demonstrate a linear scaling of the mean-squared displacement (MSD) for particles moving along the active network (Fig. S7a), consistent with long-range diffusive behavior (Fig. S7b). The effective diffusivity can be estimated from the slope of the MSD over time. Notably, the effective diffusivity of particles is highest at an intermediate value of the flow switching time $\tau$ (Fig. S7c). Flows that switch too quickly result in particles retracing their path many times along a single edge, resulting in little net effect on the diffusivity. Alternately, flows that persist too long tend to trap particles at nodes with converging flows, also decreasing their rate of spread over long distances. The existence of an optimal switching timescale was previously noted in the context of mean first passage times on active networks [34].

The optimum switching time for particle spreading is one that allows the flow directionality to persist long enough for the particle to encounter a 'trap' with converging flow, but not much longer. For a network with primarily degree-3 nodes, such as described here, a particle will cross on average 4 edges prior to encountering a trap. For the rapid velocities considered, this would correspond to an optimal switching time of $\tau^{\text{opt}} \approx 0.2$sec, corresponding to the time-scale of transition between traps. Our simulations show a peak in the effective diffusivity at switching times comparable to $\tau^{\text{opt}}$ (Fig. S7c). Notably, the optimal switching time $\tau^{\text{opt}}$ for rapid transport on the network is similar to the value ($\tau \approx 0.08$s) that allows for the most rapid local $Ca^{2+}$ release. The persistence time maximizing $Ca^{2+}$ flux is roughly half as long, presumably due to the advantage of rapid flipping in short-range delivery of $Ca^{2+}$ from the edges directly connected to the release site.

Because luminal particles are subject to both diffusion and drift, the active flows have a large effect on the long-range transport of particles with low diffusivity ($D = 3\mu\text{m}^2/\text{s}$, comparable to luminal proteins) and almost no effect on particles with high diffusivity ($D = 3\mu\text{m}^2/\text{s}$, comparable to free ions in the lumen). Thus, the presence of active network flows makes it all the more important for buffer proteins carrying $Ca^{2+}$ ions to be mobile in the lumen rather than tethered to the ER membrane.

**Network structure comparison with different release region size**

In the main text, we assume that $Ca^{2+}$ release occurs within a spatial region of radius $0.25\mu$m centered at a selected point on the ER network. This assumption is based on published data indicating that the $Ca^{2+}$ releasing region in puff events is diffraction-limited [12, 35]. In our experimental quantification of $Ca^{2+}$ puffs, the size of the releasing region cannot be resolved, due to both spatial resolution limits and the fact that the cytoplasmic $Ca^{2+}$ is expected to have spread over micro-sized distances even within the first visualized frame (3 Hz frame rate). We therefore repeat our calculations of predicted $Ca^{2+}$ release in simulations where the release site has greater spatial extent. As shown in Fig. S8, assuming a larger permeable region still results in a statistically significant difference in the cumulative release of $Ca^{2+}$ from WT versus ATL KO networks. This difference arises from both the decreased density of tubules within the release zone and the longer distances between junctions that slow down transport from distant regions of the ATL KO networks.

By contrast, for the 'bubbled ' RTN3 OE structures, the release region size has little effect on the $Ca^{2+}$ flux until the region becomes large enough to encompass multiple bubbles. At that point, the larger reservoir of $Ca^{2+}$ ions available for immediate release leads to a substantial increase in the predicted flux for the RTN3 OE structure. Even in this case, however, the release from the RTN3 OE networks does not exceed that from WT networks.

**Idealized network structures**

To highlight the importance of overall network architecture in governing the magnitude of local $Ca^{2+}$ release, we repeat our calculations on stereotypical, idealized network structures that approximately represent the ER morphology in WT and mutant cells.

WT ER forms a well-connected lattice-like network, for which the amount of tube length accessible within a given graph distance scales as would be expected for an effectively 2D system (Fig. 4f, inset). We represent this structure as a honeycomb lattice (Fig. S9a.i) with regularly spaced degree-3 nodes separated by straight edges. The edge length is taken to be $\ell \approx 0.83\mu$m, the mean edge length value found in all our extracted WT network regions.

The ER networks in ATL KO cells also behave as effectively 2D structures (Fig. 4f, inset), albeit with a decreased density and extended network tubules. To represent this morphology, we first compute the polygons formed by the edges in our extracted networks from ATL KO cells. We then calculate the aspect ratio of each polygon, defined as the longest over the shortest dimension of the bounding rectangle aligned along the principal axes of the polygon. The median aspect ratio among all extracted polygons is 2.3. We therefore took the honeycomb network used to represent WT cells and stretched it by a factor of 2.3 in one dimension, yielding an asymmetric lattice structure (Fig. S9a.ii).

For the partially fragmented ER in RTN3 OE cells, we computed the median fragment radius ($r_{\text{avg}} \approx 0.23\mu$m) and the median separation between neighboring fragments in our extracted structures ($s_{\text{avg}} \approx 1.02\mu$m). We then constructed a triangular lattice of spherical bubbles with the appropriate radius and center-to-center distances (Fig. S9a.iii). As with our extracted networks, the bubbles are treated as well-mixed spherical reservoirs, connected by narrow tubes of radius

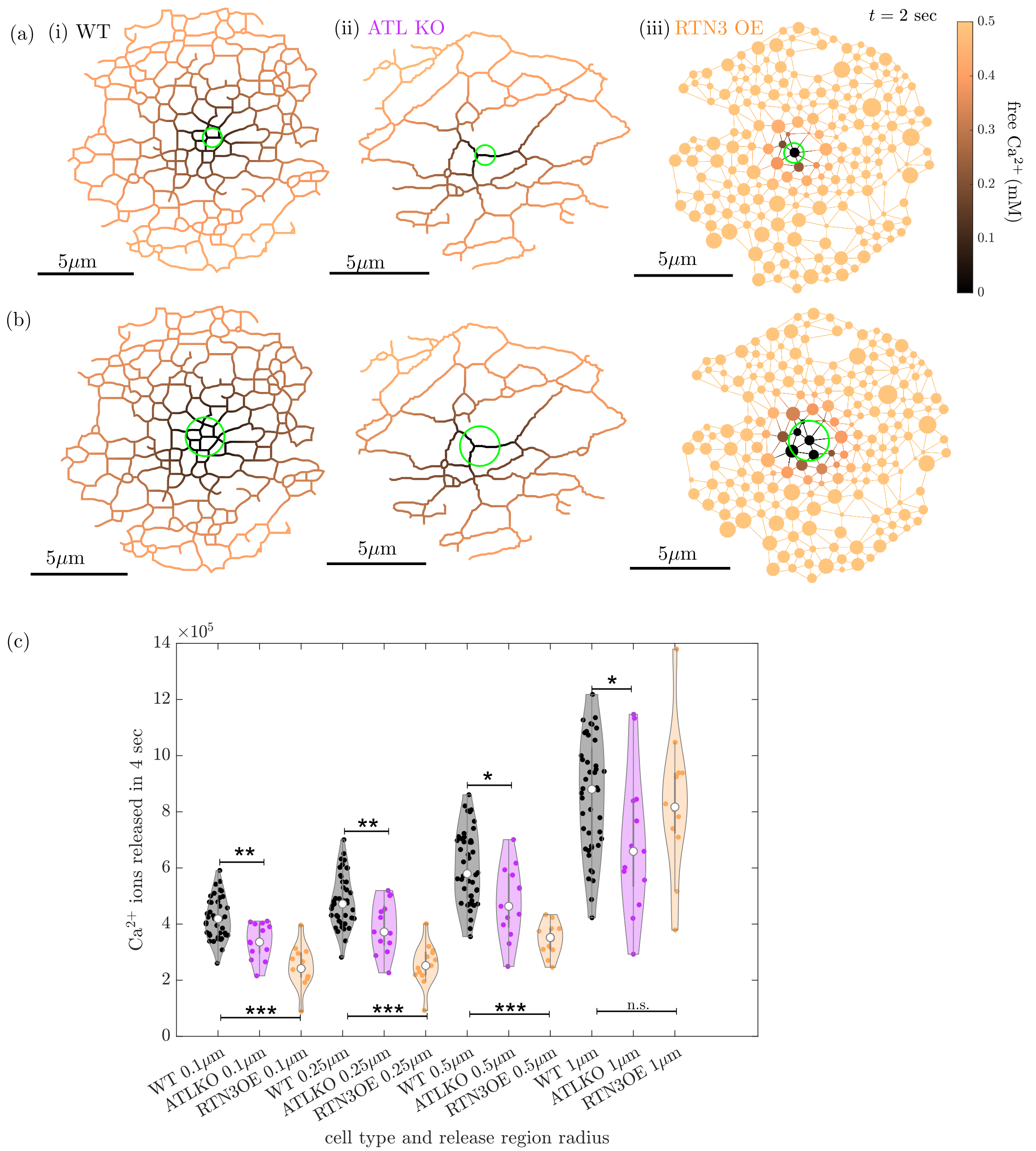


FIG. S8. $Ca^{2+}$ release from different network structures with varying release region size. (a) Simulation snapshots for extracted (i) WT, (ii) ATL KO, and (iii) RTN3 OE ER network structures, showing luminal free $Ca^{2+}$ at time 2sec after release. Release region has radius $R = 0.5\mu$m (green circle). (b) Analogous simulation snapshots with release regions of radius $R = 1\mu$m. (c) Total $Ca^{2+}$ ions released by 5sec after initiation. Each dot correponds to result for a single circular network region (as in Fig. 4). p-values computed by 2-sample Kolmogorov-Smirnov test. * $p < 0.05$, ** $p < 0.01$, *** $p < 0.001$.

10nm, with the flux from sphere into tubes governed by narrow escape dynamics (see Methods for details).

As show in Fig. S9b, the amount of $Ca^{2+}$ released over time from these idealized networks qualitatively parallels the results from extracted network structures (Fig. 4c). The increased tube length in the ATL KO network yields a slower release than WT. The poor connectivity of the

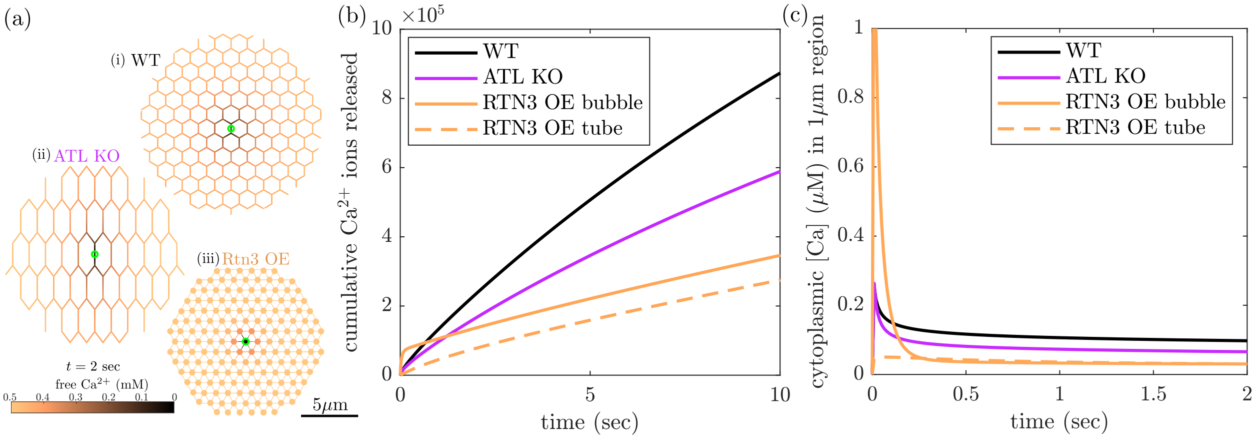


FIG. S9. Simulated $Ca^{2+}$ release from idealized network structures shows similar dynamics to extracted networks. (a) Snapshots at $t = 2$ sec after beginning of release, for 3 idealized regular network structures: (i) honeycomb network representing *WT* ER, (ii) extended honeycomb representing ATL KO ER, (iii) lattice of bubbles representing RTN3 OE ER. (b) Cumulative $Ca^{2+}$ ions released over time in the different network structures. Solid orange line shows release from a region centered on a bubbled fragment; dashed orange line shows release from a region centered on a tubule between bubbles. (c) Estimated cytoplasmic $Ca^{2+}$ levels over time, due to release from different structures as in (a) and (b).

bubbles in the RTN3 OE structure results in much slowed release after the initial transient period during which the bubble empties out. Release from the center of a tubule is lower still, due to the lack of this initial transient. Similarly, the amount of $Ca^{2+}$ expected in the cytoplasm after the first short transient period is lower in the idealized networks representing mutant cells as compared to WT (Fig. S9c). These results demonstrate that the different predicted release magnitudes for the complex heterogeneous ER networks in real cells can be largely explained by the essential changes in structure from a highly-connected lattice in WT cells, to a sparser extended lattice in ATL KO cells, and a partially fragmented poorly-connected morphology in RTN3 OE cells. The heterogeneity of real network structures accounts for the broad spread in release magnitudes (Fig. 4e) compared to the single idealized networks shown here.

##### Accounting for altered ER luminal volume

In the main text, when predicting the magnitude of $Ca^{2+}$ release from different network structures we assume a constant initial free $Ca^{2+}$ concentration ($U_0 = 0.5$mM) and a constant concentration of buffer sites $S = 2.7$mM. However, the perturbation of ER morphology is expected to alter the total volume of ER lumen, and thus potentially the $Ca^{2+}$ and buffer site concentrations. We estimate the typical ER volume in WT, ATL KO, and RTN3 OE cells as follows. First, we take the extracted network regions and compute the luminal volume per projected cell area. For WT and ATL KO cells, this value is given by $\beta = L_{\text{tot}} r^2 / R_{\text{net}}^2$ where $L$ is the total network edge length, $r = 0.05\mu$m is the tube radius, and $R_{\text{net}} = 7\mu$m is the radius of the circular domain filled by the network. For RTN3 OE cells, it is given by $\beta_{\text{RTN3}} = \left(L_{\text{tot}} r^2 + \frac{4}{3} \sum b_i^3\right) / R_{\text{net}}^2$ where $r = 0.01\mu$m for the narrow connecting tubules and $b_i$ are the radii of the bubbles. Next, we assume all cells have a perinuclear ER volume corresponding to a single sheet of height $h = 0.06\mu$m surrounding a spherical nucleus of radius $R_{\text{nuc}} = 8\mu$m, and the projected area of the cell is a circle of radius $R_{\text{cell}} = 20\mu$m. The total

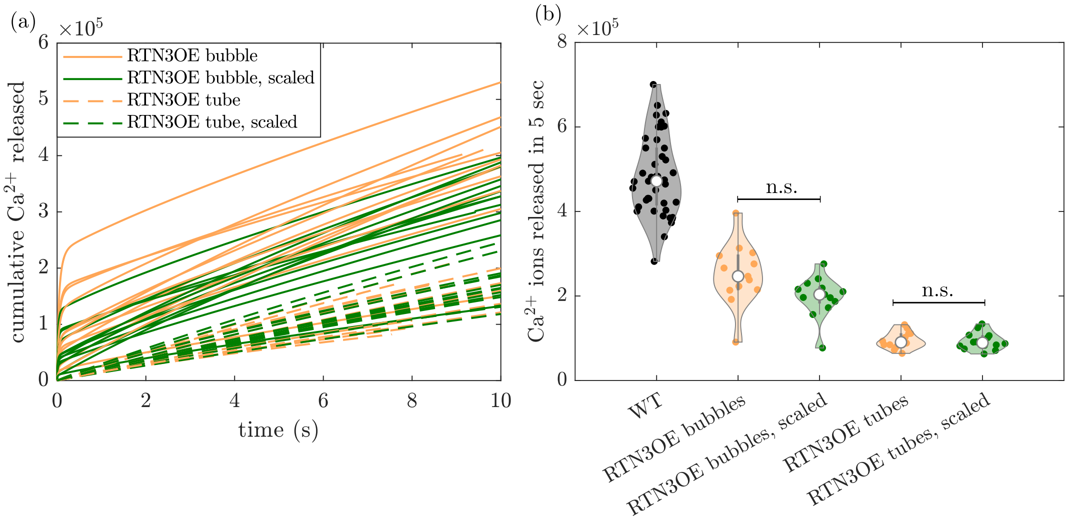


FIG. S10. Scaling concentrations allow for same total $Ca^{2+}$ load in the higher-volume RTN3 OE networks has little effect. (a) Cumulative number of $Ca^{2+}$ ions released over time from permeable regions centered on bubbles (solid lines) and tubules (dashed lines), on $n = 13$ network structure. Orange: $U_0 = 0.5$mM, $S = 2.7$mM as in Fig. 4; green: $U_0 = 0.6$mM, $S = 1.0$mM, giving same total estimated $Ca^{2+}$ load in full cell. (b) $Ca^{2+}$ ions released within first 5 sec, showing distributions for WT networks (black), and RTN3 OE networks (orange and green) with different concentrations as in (a). n.s.: not statistically significant, $p > 0.05$ via 2-sample Kolmogorov-Smirnov test.

ER volume is then estimated as

$$V_{\text{tot}} = 4\pi h R_{\text{nuc}}^2 + \beta\pi(R_{\text{circ}}^2 - R_{\text{nuc}}^2).$$

Averaging over multiple network regions and multiple cells (as in Fig. 4) gives the estimated volume for WT cells: $V_{\text{WT}} \approx 59\mu\text{m}^3$, for ATL KO cells: $V_{\text{ATL}} \approx 57\mu\text{m}^3$, and for RTN3 OE cells, $V_{\text{RTN3}} \approx 120\mu\text{m}^3$. Notably, Atlastin knockout does not appear to substantially alter the total luminal volume of the ER. However, the enlarged bubbled fragments observed in RTN3 cells lead to approximately a 2-fold increase in total volume.

Our FLIM measurements (Fig. 5) indicate that the free $Ca^{2+}$ concentration undergoes little change between the different cell types, with $U_0 \approx 0.5$mM in WT cells, and $U_0 \approx 0.6$mM in RTN3 OE and ATL KO cells. If the buffer protein concentration also stayed constant, we would expect a higher total load of $Ca^{2+}$ in the expanded volume of ER within the RTN3 OE cells. However, our thapsigargin release measurements (Fig. 5d) indicate that the total $Ca^{2+}$ load remains approximately the same in the different cell types. To account for this, we can postulate that the buffer protein production and/or import is not increased to fill the larger ER volume to the same concentration as in WT cells, and the actual concentration of buffer sites in RTN3 OE cells must be lower.

We therefore repeat our simulations in RTN3OE bubbled networks, by directly incorporating these altered concentrations. We assume for RTN3 OE cells that the initial free $Ca^{2+}$ concentration is $U_0 = 0.6$mM and the total buffer site concentration is $S = 1.0$mM, computed via Eq. S12 to give the same total load of $Ca^{2+}$ ions in the expanded volume of these cells as in the WT cell. As shown in Fig. S10a, the decreased buffer concentration is expected to slightly lower the $Ca^{2+}$ release. However, the difference is not statistically significant, considering the large variation in release magnitude due to different network structures (Fig. S10b). It should be noted that our estimates for the decreased buffer concentration consider the change in total ER volume across the entire cell. Because the bubbles of RTN3 OE networks are much larger than the tubules of WT networks, it is expected that these structures contain a greater proportion of their total $Ca^{2+}$ in the periphery. The fact that smaller releases are predicted compared to WT networks highlights the importance of transport through the network structure over and above the local volume of $Ca^{2+}$ available for immediate release.

**Effect of peripheral sheets on $Ca^{2+}$ release**

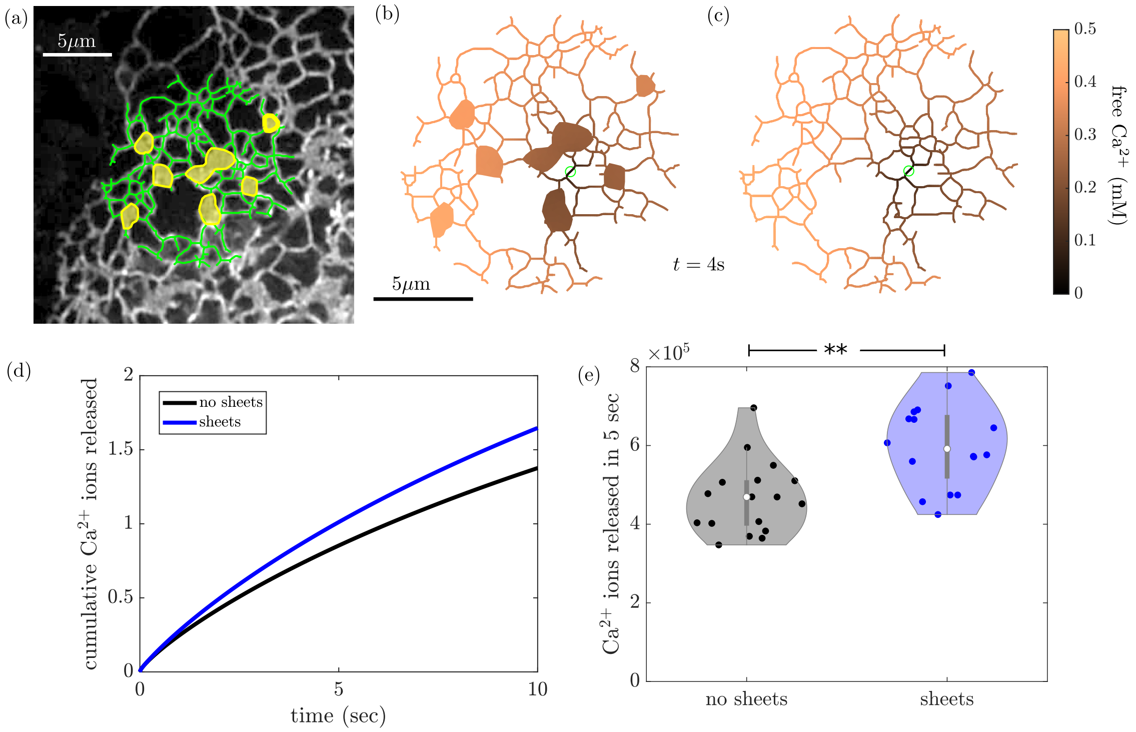


FIG. S11. Nearby peripheral sheet regions enhance $Ca^{2+}$ release. (a) Example network structure from a COS-7 cell whose peripheral ER includes sheet-like regions. Manually extracted sheet regions are marked in yellow and extracted network tubules in green. (b, c) Luminal free $Ca^{2+}$ for the extracted network, at time $t = 5$s after initiation of release. Green circle indicates the permeable region. (b) includes sheet regions in the network, while (c) fills in the corresponding regions with tubules. (d) Comparison of $Ca^{2+}$ ions released over time for the two example networks shown in (b) and (c). (e) Total $Ca^{2+}$ ions released by 4sec after initiation. Each dot marks a single network region ($n = 16$ regions from 4 different cells). The same network regions are compared with sheets present (blue) and with sheets replaced by interconnected tubules (black). ** $p \approx 0.001$ by a two-sample Kolmogorov-Smirnov test for samples being drawn from different distributions.

We briefly estimate the effect of peripheral ER sheets on the release of luminal $Ca^{2+}$. The structure of peripheral sheets remains somewhat controversial, with some high-resolution measurements indicating that apparent peripheral sheets may actually be a dense matrix of tubules and junctions [36, 37], while other observations point to the existence of dynamic nanoscale fenestrations within the sheets [11]. Here, we represent peripheral sheets with the simplest possible model – treating them as flat cisternae with a luminal thickness of 60nm (within the reported range for peripheral sheet thickness [11]).

We expand our finite volume model to include a two-dimensional triagonal meshing of sheet regions, connected to the one-dimensional meshed tubules. Sheet regions are selected manually from images of the peripheral ER in WT COS-7 cells that happen to exhibit pronounced peripheral sheets. For each network region ($14\mu$m in diameter), a release spot is selected along a tubule that is between $0.5\mu$m and $1\mu$m graph distance from the boundary of the nearest sheet. All mesh cells within $0.25\mu$m radius of the release center are marked as permeable with permeability $P = 20\mu$m/s. Fig. S11 shows the simulated release of $Ca^{2+}$ from a network interspersed with sheets. The amount of $Ca^{2+}$ released is somewhat higher than the identical comparison network where the sheets have been replaced by interconnected tubules (Fig. S11d). The simulations were repeated on 16 distinct extracted network regions from 4 different cells, again comparing the networks with sheets included to those where the sheets have been replaced by tubules. As expected, the presence of nearby sheet regions increases the predicted magnitude of $Ca^{2+}$ release (Fig. S11e) by providing a larger volume of ER lumen close to the release site to supply the $Ca^{2+}$ ions. However, the magnitude of the increase (approximately 25% difference in the median value) is relatively modest. This indicates that localizing larger reservoirs of $Ca^{2+}$ near the release region does enhance the puff magnitude, but that the effect is limited by the fact that the ions and buffer proteins still have to tunnel through the tubules to be delivered to the release site.

We next explore the effect of peripheral ER sheets on local $Ca^{2+}$ release events experimentally. The abundance of peripheral ER sheets in COS-7 cells was increased by over-expressing Climp63 (Fig. S12a) ([38–40]). We assessed the effect of increased peripheral sheets on steady state ER $Ca^{2+}$ concentration and its overall storage capacity compared to WT cells. Using the FLIM method described in the main text, we measured mTurquoise lifetime in our luminal ER $Ca^{2+}$ probe in COS-7 WT and Climp63 OE cells and did not notice any significant difference between both conditions (Fig. S12b, c). This indicates that both cells have an ER free $Ca^{2+}$ concentration of approximately 0.5mM. We estimate the total ER $Ca^{2+}$ load in WT and Climp63 OE cells, defined as integrated peak intensity of GCaMP3 in the cytoplasm after totally depleting ER $Ca^{2+}$ by thapsigargin treatment. No significant change in overall ER storage capacity is observed in this assay (Fig. S12d). Thus, cells with increased peripheral ER sheets retain high steady-state ER $Ca^{2+}$ concentration and overall storage capacity, comparable to cells with normally shaped ER.

We then proceed to quantify local $Ca^{2+}$ release events in WT and Climp63 OE cells to evaluate the effect of peripheral ER sheets on the ER's ability to supply $Ca^{2+}$ locally. We use the same $IP_3$ uncaging assay as in the main text to generate localized $Ca^{2+}$ releases and measure peak magnitude (Fig. S12e). The peak integrals are found to be slightly but significantly higher in Climp63 OE cells, suggesting that increasing peripheral ER sheets (and thus the locally available luminal volume) may increase the ability to supply $Ca^{2+}$ ions via local release (Fig. S12f). However, consistent with our modeling results (Fig. S11d, e), we do not see a drastic difference between the magnitude of $Ca^{2+}$ release from normal network structures compared to those with expanded peripheral ER sheets.

##### Effect of ER morphology on $IP_3$ receptors distribution and clustering

In the paper, we severely perturb ER morphology through RTN3 overexpression, resulting in the fragmentation of peripheral ER lumen or ATL double knockout, leading to extended tubules with fewer three-way junctions. We sought to investigate whether the depletion and over-expression of ER morphogens could interfere with $Ca^{2+}$ channel distribution and their clustering on the ER membrane. Notably, $Ca^{2+}$ can be released from the ER through the opening of inositol 1,4,5-triphosphate receptor-channels ($IP_3Rs$) [41, 42], and the number of $IP_3R$ opening determines whether the signalling event is a $Ca^{2+}$ blip, puff or $Ca^{2+}$ wave ([43, 44]).

We first assessed the distribution of $IP_3R$ in normally shaped ER using gene-edited HeLa cells in which endogenous $IP_3R1$ (major $IP_3R$ subtype in HeLa cells) is tagged with EGFP (EGFP-$IP_3R1$-

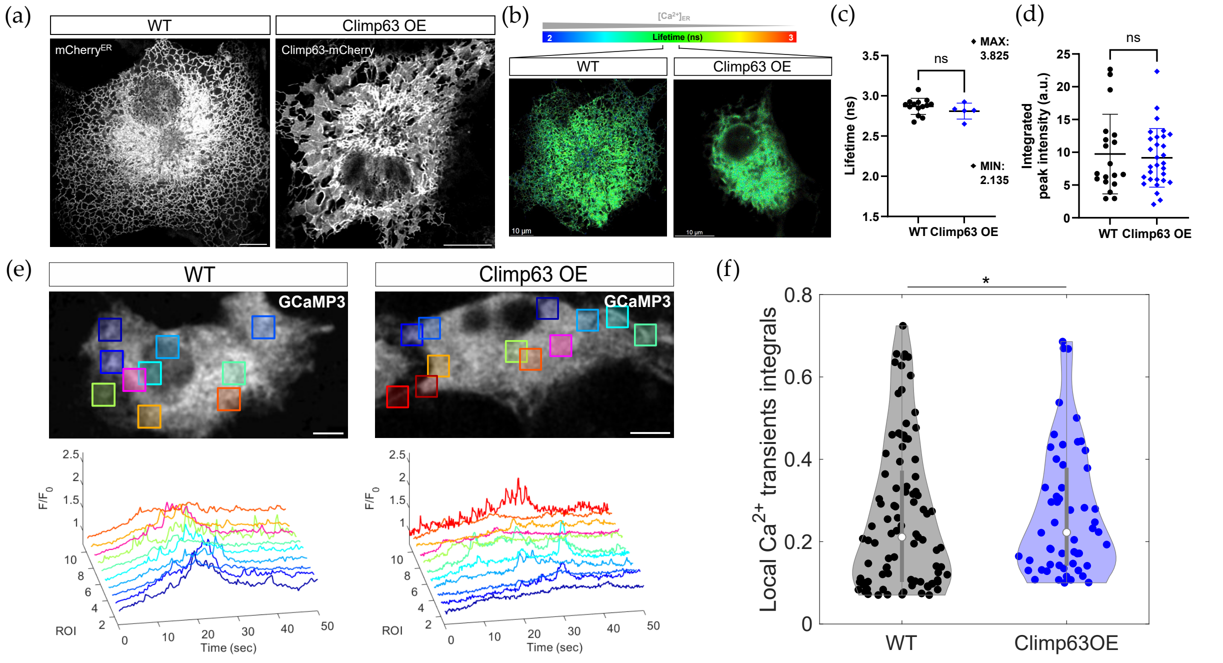


FIG. S12. Effect of increased peripheral sheets abundance on localized $Ca^{2+}$ transients and $Ca^{2+}$ storage. (a) Representative image of exogenously expressed ER luminal marker ($\text{mCherry}^{\text{ER}}$) in live COS-7 cells WT and overexpressing an ER sheet forming protein CLIMP-63 labelled with mCherry. Note the increased abundance of peripheral sheets as flat ER regions compared to the tubular peripheral ER in COS-7 WT. (b) Representative FLIM images in COS-7 cells with altered peripheral ER network structure corresponding to WT and Climp63 OE. (c) Quantitative FLIM measurements of luminal $Ca^{2+}$ concentration (each data point represents one cell). Mean lifetimes in maximal and minimal $Ca^{2+}$ conditions are indicated. (WT: $n = 14$, Climp63 OE: $n = 5$. ns: $p > 0.05$, KS test). (d) Measurement of total ER $Ca^{2+}$ load by global depletion. Integrated fluorescence intensity of GCaMP3 after addition of thapsigargin (1$\mu$M) is not significantly changed between conditions (WT: $n = 19$, Climp63 OE: $n = 30$. ns: p $>$ 0.05, KS test). (e) Representative fluorescence intensity images of GCaMP3 probe for cytoplasmic $Ca^{2+}$ released from the ER in COS-7 WT and Climp63 OE after light-induced $IP_3$ uncaging. Semi-automatically detected ROIs around local $Ca^{2+}$ transients are represented as colored squares and corresponding signal intensity traces over time within each ROI are plotted. (f) Quantification of integrated fluorescence intensity under each peak during local $Ca^{2+}$ transients (each data point represents one local $Ca^{2+}$ transient from WT: $n = 11$, Climp63 OE: $n = 13$ cells. *p $<$ 0.05, KS test). Scale bars: 10$\mu$m.

HeLa cells) ([19]). We observed the expected expression pattern of homogeneously distributed discrete puncta corresponding to several $IP_3R1$ subunits clustering together on the ER membrane in WT cells (Fig. S13a). We measured an average distance between closest neighboring clusters of 1.3$\mu$m in 3D measurements from confocal z-stack resolved with Lightning deconvolution (Fig. S13b).

We then induced ER fragmentation in EGFP-$IP_3R1$-HeLa cells by over-expressing RTN3e. $IP_3R$ clusters were also spaced by approximately 1.4$\mu$m with no significant shift in the distance separating neighboring clusters compared to WT cells (Fig. S13b). Importantly, clusters were also homogeneously distributed throughout the ER and were able to localize around inflated, seemingly vesicular portions of the ER (Fig. S13d). This confirmed that severe ER fragmentation did not prevent $IP_3R$ from localizing throughout the ER membrane and $IP_3R$ clusters were not significantly excluded from any portion of the ER network. RTN3 OE did not induce a clear rearrangement of $IP_3R$ clusters.

To determine whether $IP_3R$ clustering was affected in cells with perturbed ER morphology,

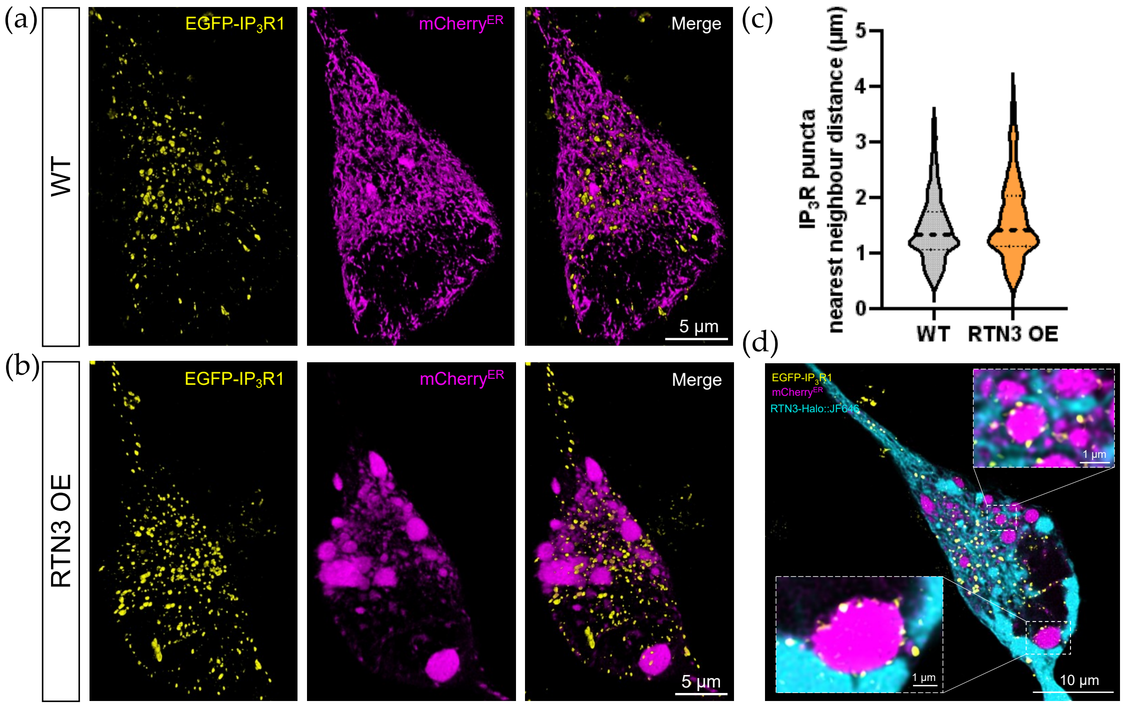


FIG. S13. $IP_3$ receptor distribution on ER tubules is maintained despite perturbation of ER architecture. (a) Representative 3D images of exogenously expressed ER luminal marker ($mCherry^{ER}$) in EGFP-$IP_3R1$ HeLa cells WT and (b) overexpressing RTN3e, from confocal 3D imaging with Lightning deconvolution. Note the punctuated expression pattern of $IP_3R1$ into discrete clusters and the fragmentation of lumen into enlarged, non-reticular structures in RTN3 OE cells in (b). (c) Quantification of the shortest distance separating two $IP_3R1$ clusters throughout the 3D z-stack (closest neighbor of the cluster center of mass, WT: $n = 2537$ puncta from $n = 12$ cells, RTN3 OE: $n = 2177$ puncta from $n = 15$ cells). (d) Representative zoom-in views of RTN3 overexpressing cell (2D slice from image in (b)) where RTN3e-Halo is labelled by JF646. Note the presence of $IP_3R1$ clusters around large ER fragments elicited by RTN3 OE. Scale bars as indicated.

we quantified the number of $IP_3R$ contained in each discrete fluorescent punctum using TIRF microscopy and single-step photobleaching. TIRF allows for the visualization of single molecules within 140nm of the plasma membrane. Images obtained from TIRF microscopy (Fig. S14a, b) confirmed that $IP_3R$ channel density was not significantly affected by ER fragmentation, with an average of 0.66 and 0.56 $IP_3R1$ per $\mu m^2$ of cell in WT and RTN3 OE cells respectively (Fig. S14e). Importantly, single-step photobleaching results (Fig. S14d) indicated that most $IP_3R1$ were found in tetramers, consistent with previous results, and this is still the case in RTN3 OE cells (Fig. S14f). Fluorescent puncta can contain several $IP_3R1$ tetramers or a single $IP_3R1$ subunit assembled into heterotetramers with other untagged subtypes of $IP_3R$, which can explain the wide distribution of the results.

Thus, severe perturbation of ER mophology did not significantly affect the homogeneous distribution of $IP_3R$ throughout the ER tubular network and its clustering into discrete puncta, likely corresponding to tetrameric channels.

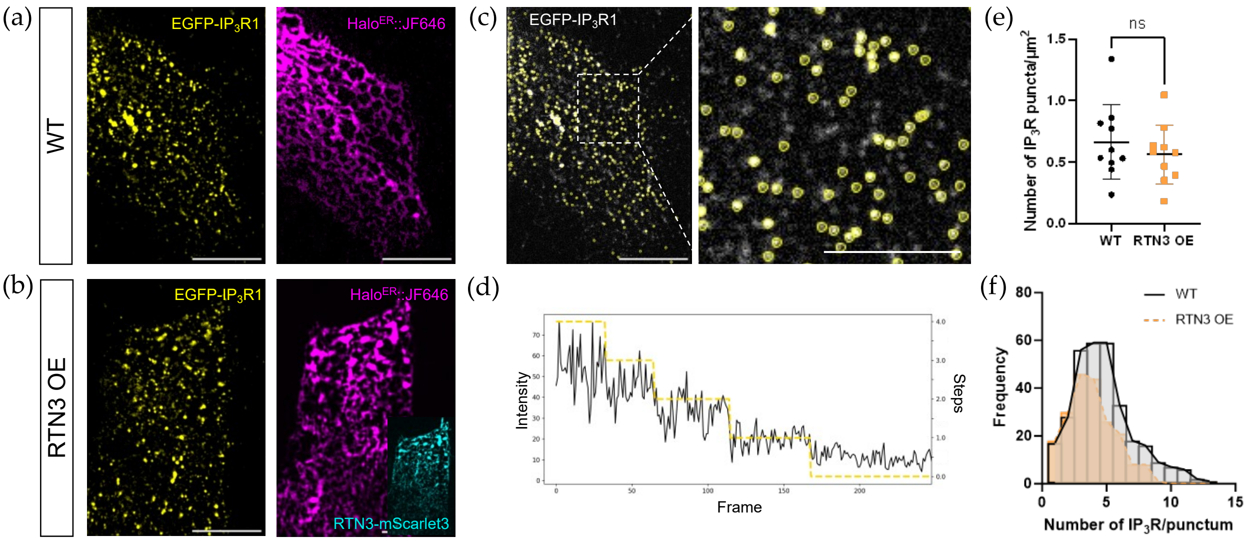


FIG. S14. $IP_3$ receptors form clusters mostly corresponding to tetramers. (a) Representative TIRF images of exogenously expressed ER luminal marker ($Halo^{ER}$ labelled with JF646) in EGFP-$IP_3$R1 HeLa cells WT and (b) overexpressing RTN3e (labelled by mScarlet3). (c) Left. Representative TIRF image of a WT EGFP-$IP_3$R1 HeLa cell where EGFP puncta are detected using TrackMate (yellow circles) with enlargement of the boxed area on the right (Scale bar: 5$\mu$m). (d) EGFP signal intensity trace of one $IP_3$R1 punctum over time showing single-step photobleaching of each fluorescent subunit in the punctum (Steps plotted as right axis, frame last 100ms). (e) Quantification of EGFP-$IP_3$R1 puncta density per cell area in TIRF images before photobleaching in WT and RTN3 OE cells (WT: $n = 10$ cells, RTN3 OE: $n = 10$ cells, ns: $p > 0.05$, KS test). (f) Frequency distribution of the number of EGFP-$IP_3$R1 per punctum from single-step photobleaching analysis (WT: $n = 320$ puncta from $n = 10$ cells, RTN3 OE: $n = 210$ puncta from $n = 10$ cells). Scale bars: 10$\mu$m.

#### S3. SUPPLEMENTAL VIDEO CAPTIONS

- **Supplemental Video 1:** Modeled release of $Ca^{2+}$ from a honeycomb network. Left: Release from passive network with purely diffusive transport of $Ca^{2+}$ ions and buffer proteins. Right: Release from active network with flows of $20\mu$m/s and persistence time $\tau_{\text{switch}} = 0.08$s. Bottom: total $Ca^{2+}$ released over time, for a single representative simulation run (average over many runs shown in Fig. 2h).

- **Supplemental Video 2:** Spatiotemporal spreading of photoactivated paGFP$^{\text{ER}}$ in the ER lumen of COS-7 WT. Representative merged images of exogenously expressed ER luminal marker (mCherry$^{\text{ER}}$ - greyscale) and paGFP$^{\text{ER}}$ signal distribution (magenta) in COS-7 cells with normal ER network (WT).

- **Supplemental Video 3:** Modeled release of $Ca^{2+}$ from three different ER network structures. Left: WT ER network. Center: ER network from ATL KO cell. Right: Fragmented ER network from RTN3 OE cell. Scale bar is $2\mu$m. Bottom: Total $Ca^{2+}$ release over time, as in Fig. 4c.

- **Supplemental Video 4:** Local $Ca^{2+}$ transients in COS-7 cells with normal ER network (WT). Representative fluorescence intensity images of GCaMP3 probe tethered in ER membrane on the cytosolic side showing cytoplasmic $Ca^{2+}$ released locally from the ER after light-induced $IP_3$ uncaging.


[1] G. D. Smith, J. Wagner, and J. Keizer, Biophys J **70**, 2527 (1996).
[2] R. Thul and M. Falcke, Biophys J **86**, 2660 (2004).
[3] S. Means, A. J. Smith, J. Shepherd, J. Shadid, J. Fowler, R. J. Wojcikiewicz, T. Mazel, G. D. Smith, and B. S. Wilson, Biophys J **91**, 537 (2006).
[4] A. J. Laude and A. W. Simpson, Febs J **276**, 1800 (2009).
[5] M. T. Alonso, M. J. Barrero, P. Michelena, E. Carnicero, I. Cuchillo, A. G. García, J. García-Sancho, M. Montero, and J. Alvarez, J Cell Biol **144**, 241 (1999).
[6] J. Meldolesi and T. Pozzan, Trends Biochem Sci **23**, 10 (1998).
[7] K.-H. Krause and M. Michalak, Cell **88**, 439 (1997).
[8] T. Konno, P. Parutto, D. M. Bailey, V. Davì, C. Crapart, M. A. Awadelkareem, C. Hockings, A. Brown, K. M. Xiang, A. Agrawal, *et al.*, bioRxiv (2021).
[9] E. A. Matthews and D. Dietrich, Front Cell Neurosci , 48 (2015).
[10] N. L. Allbritton, T. Meyer, and L. Stryer, Science **258**, 1812 (1992).
[11] L. K. Schroeder, A. E. Barentine, H. Merta, S. Schweighofer, Y. Zhang, D. Baddeley, J. Bewersdorf, and S. Bahmanyar, J Cell Biol **218**, 83 (2019).
[12] L. Bruno, G. Solovey, A. C. Ventura, S. Dargan, and S. P. Dawson, Cell Calcium **47**, 273 (2010).
[13] D. Holcman, P. Parutto, J. E. Chambers, M. Fantham, L. J. Young, S. J. Marciniak, C. F. Kaminski, D. Ron, and E. Avezov, Nat Cell Biol **20**, 1118 (2018).
[14] Z. C. Scott, K. Koning, M. Vanderwerp, L. Cohen, L. M. Westrate, and E. F. Koslover, Biophys J **122**, 3191 (2023).
[15] S. Berg, D. Kutra, T. Kroeger, C. N. Straehle, B. X. Kausler, C. Haubold, M. Schiegg, J. Ales, T. Beier, M. Rudy, *et al.*, Nat Methods **16**, 1226 (2019).
[16] W. H. Hundsdorfer, J. G. Verwer, and W. Hundsdorfer, *Numerical solution of time-dependent advection-diffusion-reaction equations*, Vol. 33 (Springer, 2003).
[17] Z. Yang and E. F. Koslover, Phys Biol (2023).

[18] O. Bénichou, C. Chevalier, J. Klafter, B. Meyer, and R. Voituriez, Nat Chem **2**, 472 (2010).
[19] N. B. Thillaiappan, A. P. Chavda, S. C. Tovey, D. L. Prole, and C. W. Taylor, Nat Commun **8** (2017), 10.1038/S41467-017-01644-8.
[20] E. Greotti, A. Wong, T. Pozzan, D. Pendin, and P. Pizzo, Sensors (Switzerland) **16** (2016), 10.3390/s16091419.
[21] S. Mehta, N. N. Aye-Han, A. Ganesan, L. Oldach, K. Gorshkov, and J. Zhang, elife **3**, 1 (2014).
[22] I. F. Smith and I. Parker, P Natl Acad Sci **106**, 6404 (2009).
[23] I. F. Smith, S. M. Wiltgen, and I. Parker, Cell Calcium **45**, 65 (2009).
[24] Z. Ozturk, C. J. O'Kane, and J. J. Pérez-Moreno, Front Neurosci **14** (2020), 10.3389/fnins.2020.00048.
[25] T. M. Inc., "Matlab version: 9.13.0 (r2022b)," (2022).
[26] K. Chen, B. Wang, J. Guan, and S. Granick, Acs Nano **7**, 8634 (2013).
[27] J. Hummert, K. Yserentant, T. Fink, J. Euchner, Y. X. Ho, S. A. Tashev, and D. P. Herten, Mol Biol Cell **32** (2021), 10.1091/MBC.E20-09-0568/ASSET/IMAGES/LARGE/MBC-32-AR35-G005.JPEG.
[28] W. Radding, S. E. Jordan, R. B. Hester, and H. C. Blair, Exp Cell Res **253**, 689 (1999).
[29] M. J. Dayel, E. F. Hom, and A. S. Verkman, Biophys J **76**, 2843 (1999).
[30] J. E. Chambers, M. Kubánková, R. G. Huber, I. López-Duarte, E. Avezov, P. J. Bond, S. J. Marciniak, and M. K. Kuimova, Acs Nano **12**, 4398 (2018).
[31] L. Xiang, R. Yan, K. Chen, W. Li, and K. Xu, Nano Lett **23**, 1711 (2023).
[32] S. Redner, *A guide to first-passage processes* (Cambridge University Press, 2001).
[33] Y. Sun, Z. Yu, C. J. Obara, K. Mittal, J. Lippincott-Schwartz, and E. F. Koslover, Physical Review Research **4**, 023182 (2022).
[34] M. Dora and D. Holcman, Proc Royal Soc B **287**, 20200493 (2020).
[35] X.-P. Sun, N. Callamaras, J. S. Marchant, and I. Parker, J Physiol **509**, 67 (1998).
[36] J. Nixon-Abell, C. J. Obara, A. V. Weigel, D. Li, W. R. Legant, C. S. Xu, H. A. Pasolli, K. Harvey, H. F. Hess, E. Betzig, *et al.*, Science **354**, aaf3928 (2016).
[37] C. J. Obara, A. S. Moore, and J. Lippincott-Schwartz, Cold Spring Harb Perspect Biol , a041259 (2022).
[38] Y. Shibata, T. Shemesh, W. A. Prinz, A. F. Palazzo, M. M. Kozlov, and T. A. Rapoport, Cell **143**, 774 (2010).
[39] G. K. Voeltz, W. A. Prinz, Y. Shibata, J. M. Rist, and T. A. Rapoport, Cell **124**, 573 (2006).
[40] Y. Shibata, C. Voss, J. M. Rist, J. Hu, T. A. Rapoport, W. A. Prinz, and G. K. Voeltz, J Biol Chem **283**, 18892 (2008).
[41] M. J. Berridge, J Exp Biol **200**, 315 (1997).
[42] M. Bootman, E. Niggli, M. Berridge, and P. Lipp, J Physiol **499**, 307 (1997).
[43] I. Parker, J. Choi, and Y. Yao, Cell Calcium **20**, 105 (1996).
[44] Y. Yao, J. Choi, and I. Parker, J Physiol **482**, 533 (1995).
